# Activity-dependent synapse elimination requires caspase-3 activation

**DOI:** 10.1101/2024.08.02.606316

**Authors:** Zhou Yu, Andrian Gutu, Namsoo Kim, Erin K. O’Shea

## Abstract

During brain development, synapses are initially formed in excess and are later eliminated in an activity-dependent manner. Weak synapses are preferentially removed, but the mechanism linking neuronal activity to synapse removal is unclear. Here we show that, in the developing mouse visual pathway, inhibiting synaptic transmission induces postsynaptic activation of caspase-3. Caspase-3 deficiency results in defects in synapse elimination driven by both spontaneous and experience-dependent neural activity. Notably, caspase-3 deficiency blocks activity-dependent synapse elimination, as evidenced by reduced engulfment of inactive synapses by microglia. Furthermore, in a mouse model of Alzheimer’s disease, caspase-3 deficiency protects against synapse loss induced by amyloid-β deposition. Our results reveal caspase-3 activation as a key step in activity-dependent synapse elimination during development and synapse loss in neurodegeneration.

## Main Text

Synapse elimination is the process in which excess synapses formed early during brain development are subsequently eliminated to form mature circuits (*1–3*). The outcome of synapse elimination depends on both spontaneous and experience-driven neural activity, with less active synapses being preferentially removed (*2, 4, 5*). However, the molecular mechanisms that connect neuronal activity to synapse elimination remain unclear. Previous studies have proposed that microglia and astrocytes, two major glial cell types of the brain, mediate activity-dependent synapse elimination by engulfing weak synapses (*6–14*), but it is unclear how glia can detect the “strength” of a synapse. A recent study proposed a molecular mechanism for activity-dependent synapse elimination in which the JAK2-STAT1 pathway is activated in inactive callosal projection neurons and regulates the removal of synapses and axons (*15*). However, the JAK-STAT pathway canonically functions through transcriptional regulation (*16*), which affects entire cells. If not all synapses made by a neuron are destined for removal (*3*), activation of the JAK2-STAT1 pathway alone may not provide sufficient elimination specificity. Therefore, other unidentified, locally-effective mechanisms likely exist that confer elimination specificity at the synapse level.

Caspase-3 is a protease crucial for the execution of apoptosis (*17*). In hippocampal neurons, transient activation of caspase-3 is required for long-term depression (LTD), a form of synaptic plasticity that induces long-lasting decreases in synaptic strength (*18*). Caspase-3 activation in apoptotic cells is also known to lead to the clearance of these cells by phagocytes (*19, 20*). These findings led us to hypothesize that caspase-3 may link synapse weakening with synapse removal by glia. Although caspases were previously implicated in dendrite pruning in metamorphosis, degeneration-like axon elimination, and the regulation of hippocampal spine density (*21–24*), the role of caspase-3 in synapse elimination in response to neural activity was not clear. In this work, we identify caspase-3 as a key molecule that is activated in postnatal development in dendritic compartments upon synaptic weakening and is necessary for elimination of weak synapses by microglia. Furthermore, we discovered that caspase-3 deficiency protects against amyloid-β-induced synapse loss in a mouse model of Alzheimer’s disease, highlighting its significance not only in developmental synapse elimination but also in adult neurodegenerative diseases.

### Tetanus toxin expression inactivates retinogeniculate synapses

We chose the mouse retinogeniculate visual pathway as a model system to study synapse elimination (*25*). In mice, retinal ganglion cells (RGCs) in each eye send out axons to form retinogeniculate synapses with relay neurons in the dorsal lateral geniculate nuclei (dLGN) of the thalamus on both sides of the brain (Fig. S1) (*25*). The majority of RGC axons from each eye innervate the contralateral (on the side opposite to the originating eye) dLGN while a smaller fraction innervate the ipsilateral (on the same side as the originating eye) dLGN (Fig. S1) (*25*). Regions in each dLGN innervated by the two eyes initially overlap but later segregate into eye-specific territories (Fig. S1). This segregation process is a hallmark of retinogeniculate pathway development at the morphological level and depends on synapse elimination and spontaneous RGC activity (*25–27*).

To investigate the role of caspase-3 in activity-dependent synapse elimination, we first needed to establish a method that could manipulate the strength of a subset of retinogeniculate synapses. For this purpose, we used adeno-associated-virus (AAV) to deliver a construct expressing tetanus toxin light chain (TeTxLC) under the control of the neuron-specific human synapsin promoter (AAV-hSyn-TeTxLC) (Fig. 1A) (*28*). *In utero* intraocular injection of AAV-hSyn-TeTxLC at embryonic day 15 (E15) leads to TeTxLC expression in RGCs by the time of birth, blocking neurotransmitter release at retinogeniculate synapses by cleaving synaptobrevin and leading to synapse inactivation (*28, 29*). To label RGC axons and facilitate quantification, we co-injected AAVs that express fluorescent proteins (mTurquiose2, eGFP, or tdTomato, depending on the experiment) as anterograde tracers (Fig. 1A). These AAVs expressing fluorescent proteins also served as controls for eye injections.

**Figure 1.**
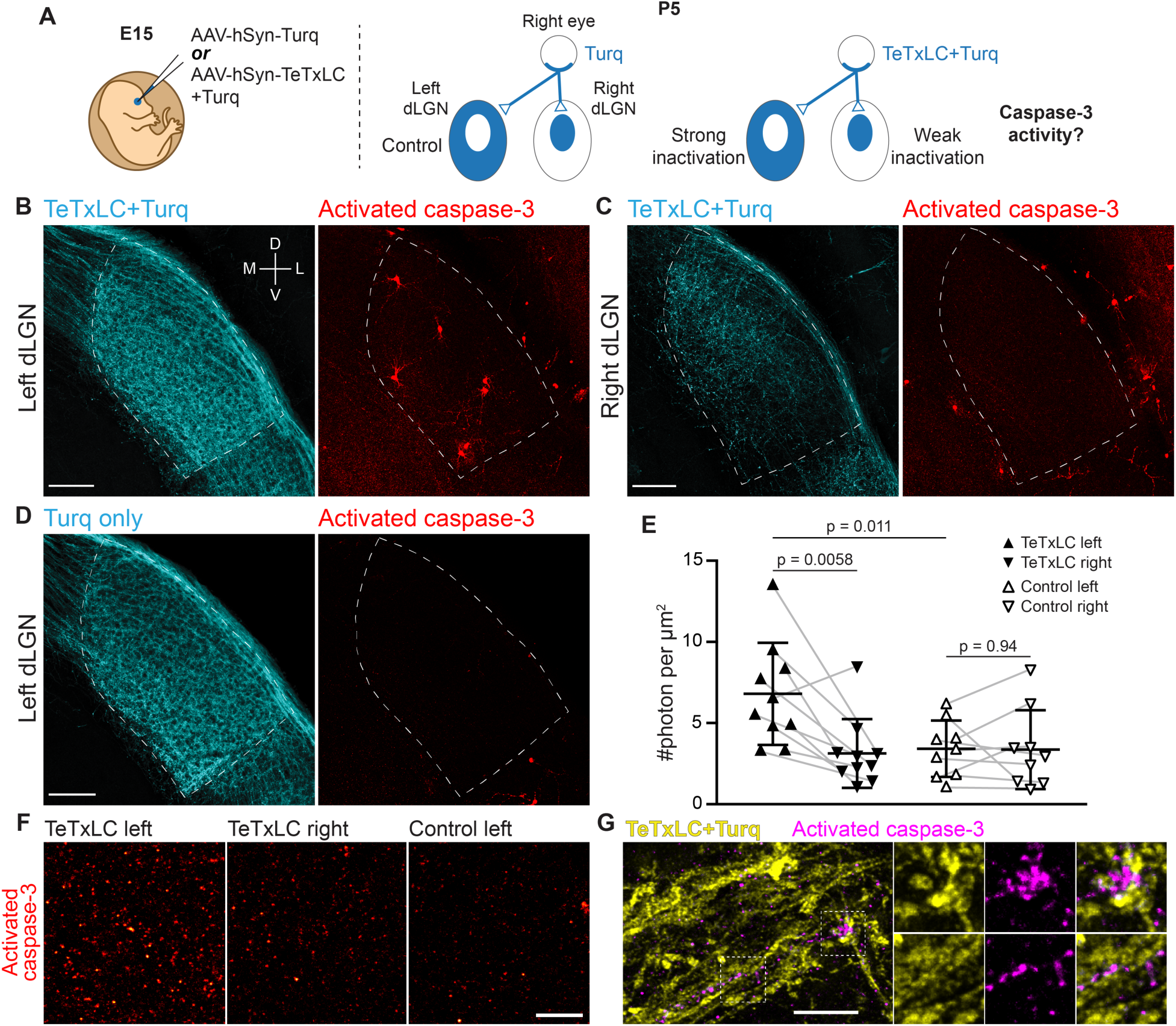
Inactivation of retinogeniculate synapses induces caspase-3 activity. (**A**) Schematics of experimental setup. AAVs expressing tetanus toxin light chain (TeTxLC) and/or mTurquiose2 (Turq) were injected into the right eye of E15 mice (left). By P5, retinogeniculate synapses in dLGN were inactivated to varying extents depending on injection and side (right). (**B**-**D**) Confocal images of Turq (left panels) and activated caspase-3 (right panels) in left dLGN (**B**) and right dLGN (**C**) of a TeTxLC-injected P5 animal and in left dLGN of a control P5 animal (**D**). Dotted lines delineate dLGN boundaries. Only signals within dLGNs were analyzed. Images from the same fluorescent channel were adjusted to the same contrast. The compass in **B** marks tissue orientation. Scale-bars: 100 μm. D, dorsal; V, ventral; M, medial; L, lateral. (**E**) Quantification of caspase-3 activity in indicated dLGNs. Activated caspase-3 signals in each dLGN (highlighted areas in **B**-**D**) were summed and normalized to dLGN area. Each point represents the result from one dLGN. Data from two dLGNs of the same animal were paired for analysis (grey lines). n=10 for TeTxLC-injected animals and n=9 for control animals. Mean and standard deviation (S.D.) are shown. P-values were calculated from two-tailed t-tests (paired when applicable). (**F**) Example images showing punctate caspase-3 activities in ventral-medial regions of indicated dLGNs. Images were adjusted to the same contrast. Scale-bar: 20 μm. (**G**) High-resolution images of dLGN showing TeTxLC-expressing RGC axons (yellow) and activated caspase-3 (magenta). Two regions of interest (dotted squares) are magnified to illustrate that caspase-3 activity was found juxtaposing TeTxLC-expressing axon terminals but not within them. Scale-bar: 5 μm.

To validate the efficacy of our synapse inactivation method, we injected AAV-hSyn-TeTxLC into the right eye of wildtype E15 embryos and analyzed the segregation of eye-specific territories at postnatal day 8 (P8), when the segregation process is largely complete (*25*). To quantify the overlap between eye-specific territories in an unbiased manner, we applied a set of increasing thresholds to signals from both eyes (Fig. S2) (*30*). For each threshold, we calculated a percentage overlap value as the ratio of the dLGN area with signals from both eyes to the total dLGN area (Fig. S2 and Fig. S3). Consistent with previous results (*26, 27*), inactivating right eye-associated retinogeniculate synapses led to contraction of right eye-specific territories and expansion of left eye-specific territories relative to control animals (Fig. S3A and S3B). Importantly, segregation of eye-specific territories in TeTxLC-injected mice was significantly defective (Fig. S3C and S3D), confirming that our method potently inactivated synapses and perturbed activity-dependent refinement of the retinogeniculate pathway.

### Synapse inactivation induces postsynaptic caspase-3 activity

Is caspase-3 activated in response to synapse weakening? To address this question, we injected AAV-hSyn-TeTxLC into the right eyes of E15 embryos and looked for caspase-3 activation in dLGNs of TeTxLC-injected mice at the age of P5 (a time when synapse elimination is most active) (Fig. 1A) (*7, 25*). We used immunohistochemistry against the cleaved and active form of caspase-3 to detect caspase-3 activity in the dLGN area (*17*). If synapse inactivation leads to caspase-3 activation, we would expect higher levels of caspase-3 activity in dLGNs with more inactivated synapses (Fig. 1A). In agreement with this hypothesis, caspase-3 activity in left dLGNs of TeTxLC-injected animals (dLGNs with the most inactivated synapses) was significantly higher compared to that in right dLGNs of the same animals (dLGNs with intermediate amount of inactivated synapses) (Fig. 1B, 1C, and 1E) and compared to that in dLGNs of animals receiving control injections in the right eyes (dLGNs with no inactivation) (Fig. 1B, 1D, and 1E). Reassuringly, caspase-3 activities in left and right dLGNs of control animals were comparable (Fig. 1E), despite injections being administered only in the right eyes. This suggests that surgical manipulation and viral transduction had no measurable contribution to caspase-3 activation. These findings support the model that inactivation of retinogeniculate synapses leads to caspase-3 activation.

Does inactivation of retinogeniculate synapses activate caspase-3 pre-synaptically in RGCs or post-synaptically in dLGN relay neurons? Caspase-3 activity in dLGNs of TeTxLC-injected mice was present either as discrete, punctate signals (Fig. 1F) or as staining of entire cells (Fig. 1B and Fig. S4Bi-iii). Both types of signals were more abundant in dLGNs with strong synapse inactivation than in control dLGNs (Fig. 1B, 1D and 1F). Intriguingly, punctate caspase-3 activity was not found within TeTxLC-expressing RGC axons (Fig. 1G). Instead, punctate caspase-3 activity closely juxtaposed inactivated axon terminals (Fig. 1G) and co-localized with the dendritic marker MAP2 (Fig. S4A). In the case where caspase-3 was activated in entire cells, the cells were positive for the neuronal marker NeuN (Fig. S4C) and possessed relatively large round somas and multipolar dendritic arbors (Fig. S4Bi-iii), all of which are characteristics of dLGN relay neurons (*31, 32*). These observations suggest that caspase-3 activation triggered by synapse inactivation occurs predominantly in the dendritic compartments of dLGN relay neurons that are postsynaptic to the inactivated synapses.

Although morphologically distinct, localized and whole-cell caspase-3 activity may be mechanistically linked. In support of this, we observed transitional states of caspase-3 activation where multiple dendrites of a relay neuron were positive for active caspase-3 without the cell body becoming apoptotic (an example is shown in Fig. S4Biv), suggesting that whole-cell caspase-3 activation may arise from accumulation of local caspase-3 activity. Such transitions might be particularly relevant in dLGNs that receive the majority of TeTxLC-expressing axons, as systematic silencing of synapses could lead to widespread caspase-3 activation in the postsynaptic neuron, potentially overwhelming negative regulatory mechanisms that normally keep caspase-3 activation localized (*24, 33–35*). While our data cannot determine which form of caspase-3 activation plays a more significant role in the normal development of the visual pathway, the infrequent occurrence of whole-cell caspase-3 activation in control dLGNs (Figure 1D) -- which undergo normal synapse elimination -- suggests that localized caspase-3 activation is necessary to drive synapse elimination under physiological conditions.

### Caspase-3 activation at weak synapses requires the presence of strong synapses

Previous experiments have manipulated RGC neuronal activities and demonstrated that, regardless of the absolute level of RGC activity, the dLGN territory of less active RGCs always contracted relative to that of more active RGCs (*26, 27*). This observation suggests that the outcome of synapse elimination is driven by competitive interactions between strong and weak synapses. Consistent with this idea, a recent study demonstrated that elimination of weak synapses in callosal projections occurred only when other strong synapses are present (*28, 36*).

To test if caspase-3 activation at weak synapses requires the presence of strong synapses, we designed two experimental conditions. In the first condition, we inactivated RGCs only in the right eye, leading to the presence of both active (from the left eye) and inactive (from the right eye) synapses in the dLGN (Fig. 2A). We termed this condition “single inactivation”. In the second condition, we inactivated RGCs in both eyes, leading to the presence of only inactive synapses in the dLGN (Fig. 2A). We termed this condition “dual inactivation”. We also included a control condition where no RGCs were inactivated (Fig. 2A). We first confirmed that, in accordance with previous findings (*26, 37*), segregation of eye-specific territories was disrupted in the dual inactivation condition (Fig. S5A-D). We then measured dLGN caspase-3 activity in control, single, and dual inactivation conditions. Intriguingly, while dLGN caspase-3 activity was significantly higher in the single inactivation condition compared to that in the control (Fig. 2B and 2C), dLGN caspase-3 activity in the dual inactivation condition was not significantly different from that in the control (Fig. 2B and 2C) and was significantly lower than that in the single inactivation condition (Fig. 2B and 2C). Although not reaching statistical significance, dLGN caspase-3 activity on average was still higher in the dual inactivation condition compared to the control (Fig. 2C), possibly due to incomplete inactivation of retinogeniculate synapses, or interaction between weak retinogeniculate synapses and strong synapses from other pathways, or additional mechanisms that activate caspase-3 independently of interactions between weak and strong synapses. These results suggest that interaction between weak and strong synapses is required for caspase-3 activation to occur at weak synapses.

**Figure 2.**
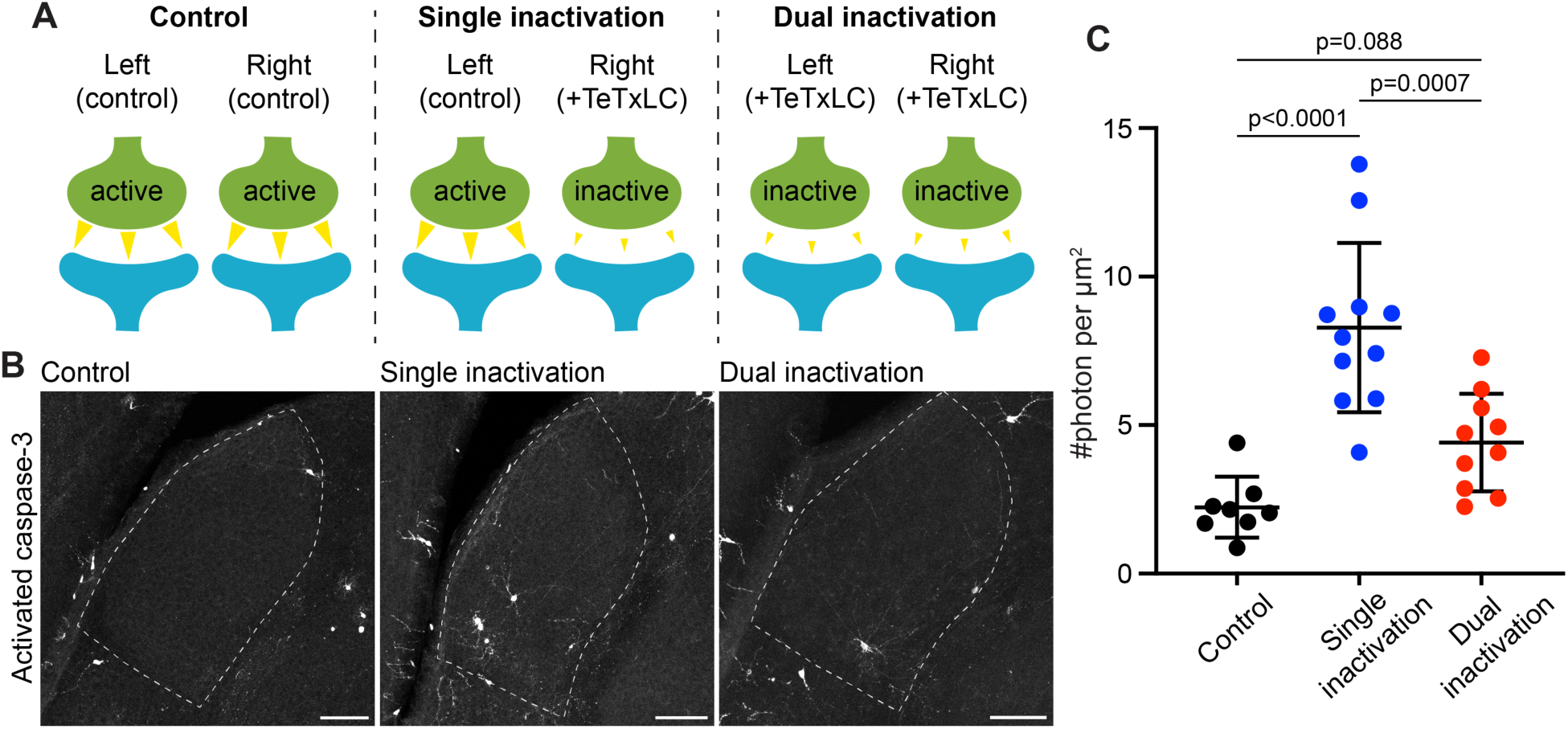
Caspase-3 activation at weak synapses requires the presence of strong synapses. (**A**) Schematics illustrating experimental conditions. No synapses are inactivated in the control (left); only synapses from right eyes are inactivated in single inactivation condition (middle); synapses from both right and left eyes are inactivated in dual inactivation condition (right). (**B**) Confocal images of P5 left side dLGNs in the three conditions showing caspase-3 activity in the dLGN. Dotted lines mark dLGN boundaries. Scale-bars: 100 μm. (**C**) Quantification of dLGN caspase-3 activity in the indicated conditions. Caspase-3 activity in each dLGN were summed and normalized to the dLGN area. For the single inactivation condition, values were from left dLGNs only. For the other two conditions, values from both dLGNs were averaged. n=8 animals for control, n=11 animals for single inactivation, and n=10 animals for dual inactivation. Mean and S.D. are shown. P-values were calculated from Tukey’s multiple comparison tests.

### Caspase-3 is required for segregation of eye-specific territories

Is synapse inactivation-induced caspase-3 activity important for synapse elimination in the retinogeniculate pathway? To address this question, we investigated whether segregation of eye-specific dLGN territories is defective if caspase-3 activation does not occur. Since we were unable to find a Cre-expressing mouse line that satisfactorily depletes caspase-3 in dLGN relay neurons, we used full-body caspase-3 deficient (*Casp3*^−/−^) mice (*38, 39*). These mice were backcrossed to congenicity on the C57Bl/6J background and were viable, fertile, and did not exhibit overt abnormalities (*39*). To measure segregation of eye-specific territories, we injected the eyes of *Casp3*^+/+^ and *Casp3*^−/−^ littermate mice at P9 with fluorophore-conjugated cholera toxin subunit B (CTB), using a different fluorophore for each eye. CTB molecules bind to ganglioside molecules on RGC surfaces and anterogradely label RGC axon terminals (*40*). At P10, when CTB labeling and eye-specific segregation were complete (*25, 30*), we collected the brains and imaged eye-specific territories in each dLGN. By visual inspection, eye-specific territories in *Casp3*^+/+^ dLGNs overlapped minimally (Fig. 3A). In contrast, *Casp3*^−/−^ dLGNs showed clear defects in the separation of eye-specific territories (Fig. 3B) (*41*). We then quantified eye-specific segregation as described previously (Figure S2) and found that percentage overlap in *Casp3*^−/−^ dLGNs was consistently higher than that in *Casp3*^+/+^ dLGNs across all thresholds (Fig. 3C and D). Importantly, the fold difference and statistical significance of the difference in percentage overlap between the two groups of animals increased as the cutoff threshold increased (Fig. 3D), suggesting the defects observed in *Casp3*^−/−^ animals cannot be attributed to artifacts generated from background noise.

**Figure 3.**
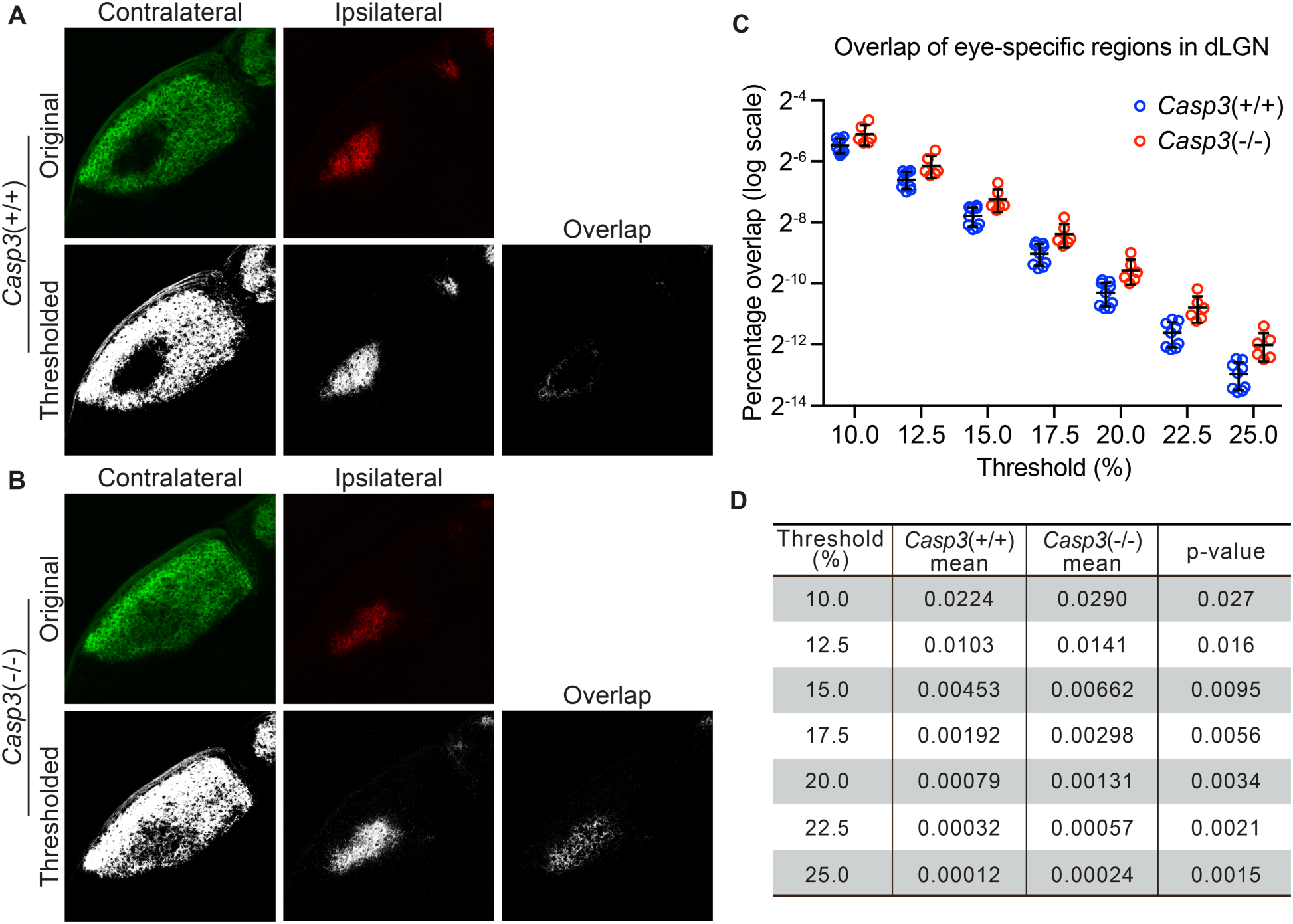
Caspase-3 is required for segregation of eye-specific territories. (**A**-**B**) Representative confocal images of retinogeniculate inputs in the dLGN of P10 wild-type (**A**) and *Casp3*^−/−^ (**B**) mice. Contralateral inputs are labeled with AlexaFlour488 (AF488) conjugated cholera toxin subunit B (CTB) and ipsilateral inputs with AF594-CTB. Original images were thresholded into 0-or-1 images using the Otsu method (*34*), and the overlap between thresholded contralateral and ipsilateral inputs is shown. (**C**) Percentage overlap between eye-specific territories in wildtype and *Casp3*^−/−^ mice under a series of increasing signal cutoff thresholds. Note that the percentage overlap is plotted on a log scale. Each circle represents one animal. Mean and S.D. are shown. n=9 for wildtype mice and n=6 for *Casp3*^−/−^ mice. (**D**) Mean percentage overlap values in wildtype and *Casp3*^−/−^ mice and p-values of two-tailed t-tests between the two genotypes are listed for each cutoff threshold.

As RGCs are known to undergo extensive apoptosis during the first week of postnatal development (*42, 43*), we wondered if caspase-3 deficiency might block RGC turnover and result in more RGC inputs in the dLGN, thereby confounding the overlap analysis. To address this concern, we quantified the density of RGCs in whole-mount retinae using a RGC marker, RBPMS (Fig. S6A) (*44*). Reassuringly, RGC densities in retinae of *Casp3*^+/+^ and *Casp3*^−/−^ mice were not different (Fig. S6B), indicating that RGC apoptosis still occurred at normal rates in the absence of caspase-3, presumably through redundant mechanisms.

It is also possible that caspase-3 deficiency might confound the overlap analysis by affecting the number of dLGN relay neurons. Therefore, we used NeuN staining to visualized relay neurons in dLGNs of *Casp3*^+/+^ and *Casp3*^−/−^ mice and confirmed that relay neuron density is not significantly altered by caspase-3 deficiency (Figure S7). Notably, this observation indicates that, during normal development, synapse weakening does not induce widespread whole-cell caspase-3 activation in dLGN relay neurons, corroborating the idea that localized caspase-3 activity drives synapse elimination in dLGN. Collectively, our findings suggest that caspase-3 is required for normal segregation of eye-specific territories in dLGN.

### Caspase-3 is required for retinogeniculate circuit refinement

Segregation of eye-specific territories in the mouse dLGN occurs prior to eye opening (which happens at ∼P13) and relies predominantly on spontaneous RGC activity (*25*). After eye opening, sensory-dependent RGC activity drives a second phase of refinement at the circuit level (*25*). At the time of eye opening, each dLGN relay neuron receives numerous weak RGC inputs. With visual experience, a minority of these inputs are strengthened while other weak inputs are eliminated, so that by P30 only a few (typically ≤3) strong RGC inputs innervate each dLGN relay neuron (*25*).

To test if caspase-3 is required for visual experience-driven circuit refinement, we prepared acute brain slices from P30 *Casp3*^+/+^ and *Casp3*^−/−^ littermate mice and investigated the electrophysiological properties of retinogeniculate synapses (Fig. S8A) (*45, 46*). We stimulated the optic tract while recording both AMPA receptor (AMPAR)- and NMDA receptor (NMDAR)-mediated excitatory postsynaptic currents (EPSCs) in dLGN relay neurons (Fig. S8A and Fig. 4A-B). By gradually increasing the stimulation intensity on the optic tract, we recruit individual RGC axons in a serial manner (Fig. S8A). Excitation of each RGC axon triggers a step increase in the EPSC amplitude (Fig. S8A), allowing us to infer RGC input numbers by counting steps in EPSC response curves (Fig. 4C and S9A). In agreement with previous studies, retinogeniculate circuits in *Casp3*^+/+^ animals were refined, with dLGN relay neurons most frequently receiving 2 strong RGC inputs, and the great majority of neurons receiving no more than 3 inputs (Fig. 4A, 4C and S9A). In contrast, RGC input counts of dLGN relay neurons in *Casp3*^−/−^ animals followed a long-tailed distribution that was significantly right-shifted from that of *Casp3*^+/+^ animals (Fig. 4C and S9A). The majority of *Casp3*^−/−^ relay neurons received more than 3 inputs (Fig. 4B-C and S9A), and many inputs elicited only small increments in EPSC amplitude (Fig. 4B). Additionally, we examined maximum EPSC amplitude (Fig. S9B-C), AMPAR-EPSC to NMDAR-EPSC ratio (Fig. S9D), and fiber fraction (Fig. S9E) (*46*) in *Casp3*^+/+^ and *Casp3*^−/−^ mice. We found no significant difference between the two groups of animals, suggesting that the defective circuit refinement in *Casp3*^−/−^ animals was not due to the lack of retinogeniculate synapse maturation but instead an inability to eliminate weak synapses.

**Figure 4.**
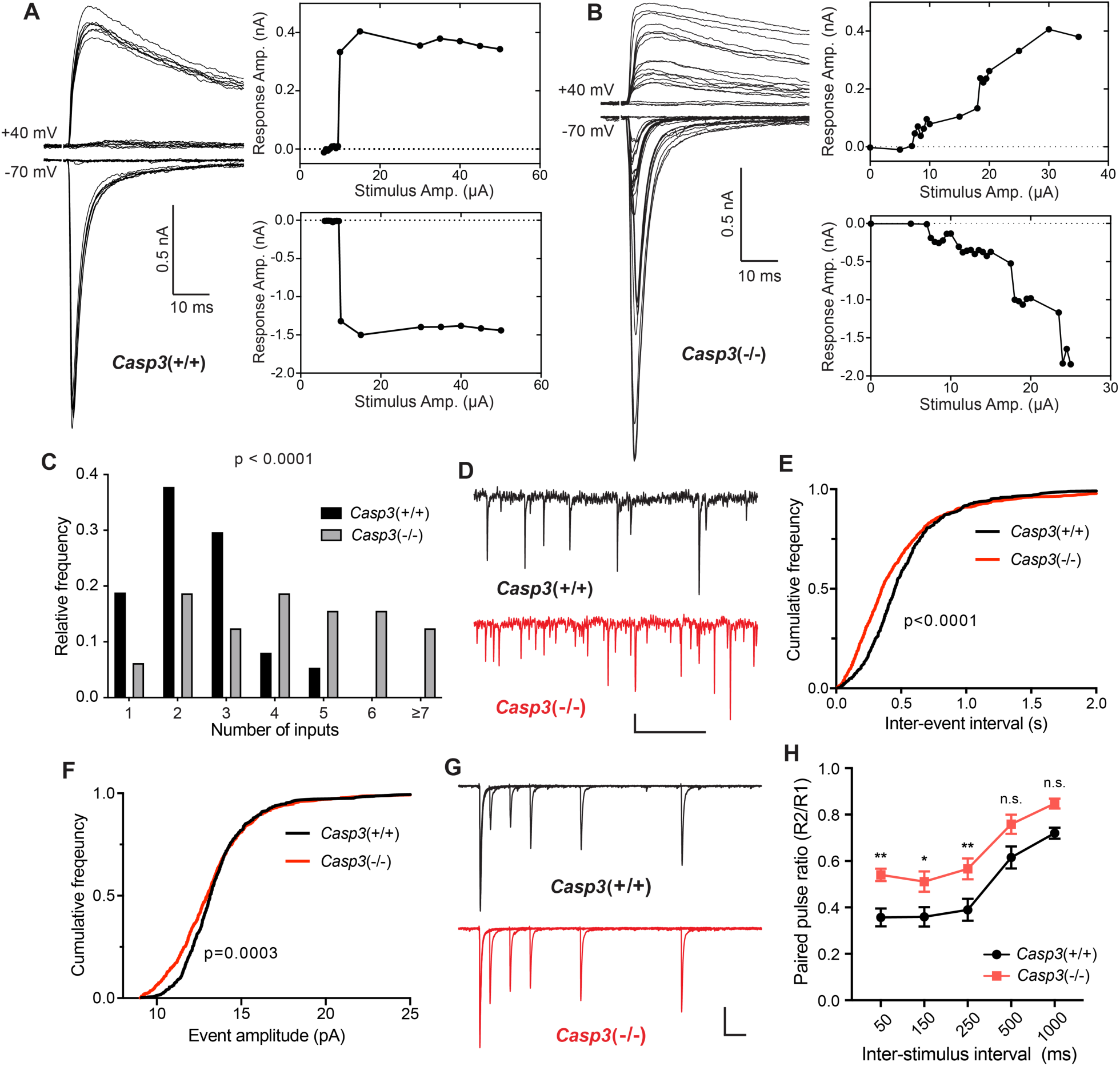
Caspase 3 is required for retinogeniculate circuit refinement. (**A**-**B**) Example recordings of dLGN relay neuron responses in P30 wildtype (**A**) and *Casp3*^−/−^ (**B**) mice. Excitatory postsynaptic currents (EPSCs) were evoked by increasing stimulation currents in the optic tract. Both AMPAR-mediated inward current at -70 mV membrane potential and AMPAR and NMDAR-mediated outward current at +40 mV membrane potential are shown. Peak response amplitudes at each stimulation intensity are plotted to the right of recording traces. Scale bars represent 0.5 nA and 10 ms. (**C**) Distribution of RGC input numbers on individual dLGN relay neurons in wildtype and *Casp3*^−/−^ mice. The number of RGC inputs was inferred by manually counting the number of steps in AMPAR-mediated EPSC response curves (lower right in **A** and **B**) while blind to the genotypes. P-value was calculated from two-tailed t-test. n=37 cells from 22 wildtype mice and n=32 cells from 16 *Casp3*^−/−^ mice. (**D**) Example recordings of mEPSC measurements from wildtype and *Casp3*^−/−^ mice. Scale-bars represent 0.5 s (horizontal) and 5 pA (vertical). (**E**-**F**) Cumulative distribution curves of inter-mEPSC intervals (**E**) and mEPSC amplitudes (**F**) in wildtype and *Casp3*^−/−^ mice. P-values were calculated from Kolmogorov-Smirnov tests. n=16 cells from 4 wildtype mice and n=17 cells from 4 *Casp3*^−/−^ mice. (**G**) Example recordings from paired pulse measurements at -70 mV membrane potential in wildtype and *Casp3*^−/−^ mice. Traces from experiments with 50, 150, 250, 500, and 1000 ms inter-stimulus intervals are overlayed. Stimulus artifacts were removed from the traces for clarity. Scale-bars represent 100 ms (horizontal) and 0.3 nA (vertical). (**H**) Paired-pulse ratio (calculated as amplitude of the second response over that of the first response) in wildtype and *Casp3*^−/−^ mice at various inter-stimulus intervals. Mean and standard error of the mean (SEM) are shown. P-values were calculated from Bonferroni’s multiple comparison test. p=0.0067 for 50 ms interval, p=0.0369 for 150 ms interval, and p=0.0097 for 250 ms interval. n=16 cells from 7 wildtype mice and n=13 cells from 4 *Casp3*^−/−^ mice.

An increased number of RGC inputs innervating dLGN relay neurons in *Casp3*^−/−^ mice should correspond to greater numbers of presynaptic release sites and should result in more frequent spontaneous release of neurotransmitters. To test this prediction, we measured miniature EPSCs (mEPSCs) in dLGN relay neurons of *Casp3*^+/+^ and *Casp3*^−/−^ mice (Fig. 4D). We found that mEPSCs in *Casp3*^−/−^ neurons occurred more frequently (Fig. 4E), with a greater fraction of them having small amplitudes (Fig. 4F). These results mirror previous measurements in hippocampal CA1 neurons in Casp3^−/−^ mice (*24*). Overall, our observations are consistent with the model where caspase-3 deficiency impairs the removal of weak synapses, leading to greater number of inputs on dLGN relay neurons.

As an additional way to probe retinogeniculate circuit refinement, we measured paired-pulse response ratios (PPRs) in dLGN relay neurons (Fig. S8B and Fig. 4G-H) (*45, 47*). The retinogeniculate circuit typically generates PPRs that are smaller than one due to depletion of neurotransmitters (Fig. S8B) (*45, 47*). If there is an increase in the number of release sites in *Casp3*^−/−^ mice, we expect more available neurotransmitters and larger PPRs (Fig. S8C) (*47*). Indeed, we observed higher PPRs in *Casp3*^−/−^ dLGN relay neurons (Fig. 4D-E), suggesting that defective synapse elimination resulted in more RGC release sites in *Casp3*^−/−^ mice. Taken together, our results demonstrate that caspase-3 is required for experience-driven retinogeniculate circuit refinement.

### Caspase-3 deficiency impairs synapse elimination by microglia

Our results demonstrate that proper synapse elimination in the retinogeniculate pathway requires caspase-3 activity. Previous studies have shown that microglia and astrocytes engulf weak synapses during the process of synapse elimination (*7, 10*). We therefore sought to determine whether caspase-3 activation is required for synapse elimination by microglia and/or astrocytes.

If synapse elimination by microglia depends on caspase-3 activation, then microglia in *Casp3*^−/−^ mice should engulf less synaptic material. Similar to a previous study (*7*), we utilized the *Cx3cr1-Gfp* transgenic mouse line to visualize microglia *in vivo.* In this line, the native *Cx3cr1* promoter drives *Gfp* expression and specifically labels microglia in the brain (Fig. 5A) (*48*). We labeled RGC axon terminals by injecting fluorophore-conjugated CTB in the eyes of P4 *Casp3*^+/−^; *Cx3cr1-Gfp*^+/−^ and *Casp3*^−/−^; *Cx3cr1-Gfp*^+/−^ littermate pups (*Casp3*^+/−^ instead of *Casp3*^+/+^ mice were used to enable more efficient breeding) (*7*). At P5, the brains were harvested and the dLGN imaged (Figure 5A). Microglia in *Casp3*^−/−^; *Cx3cr1-Gfp*^+/−^ mice had small soma and thin processes, indicating that caspase-3 deficiency did not result in microglial activation (Fig. 5B). This is further validated by immunohistochemical analysis of microglia in caspase-3 deficient mice (Fig. S10A-D). To quantify synapse engulfment, we isolated volumes corresponding to microglia and RGC axon terminals in the dLGN (Fig. 5B) and calculated the volume of RGC terminals that were fully enclosed in each microglia (Fig. 5C). As expected, we observed that microglia in *Casp3*^−/−^; *Cx3cr1-Gfp*^+/−^ mice engulfed significantly less axonal material compared to those of *Casp3*^+/−^; *Cx3cr1-Gfp*^+/−^ littermates (Fig. 5B-C). The same trend was observed when RGC axon terminals originating from the two eyes were analyzed separately (Fig. S11A-B). As the engulfment capacity of microglia is limited by their size, we normalized the volume of engulfed synaptic material in each microglia to the microglial volume (Fig. 5D). Microglia in both groups of mice are comparable in size (Fig. S11C), and the defective synaptic engulfment in *Casp3*^−/−^ mice persisted after normalization to microglial volume (Fig. 5D). Additionally, the same defect was observed when engulfment values were averaged and reported by animal (Fig. S11D). Taken together, our results demonstrate that microglia-mediated synapse elimination depends on caspase-3.

**Figure 5.**
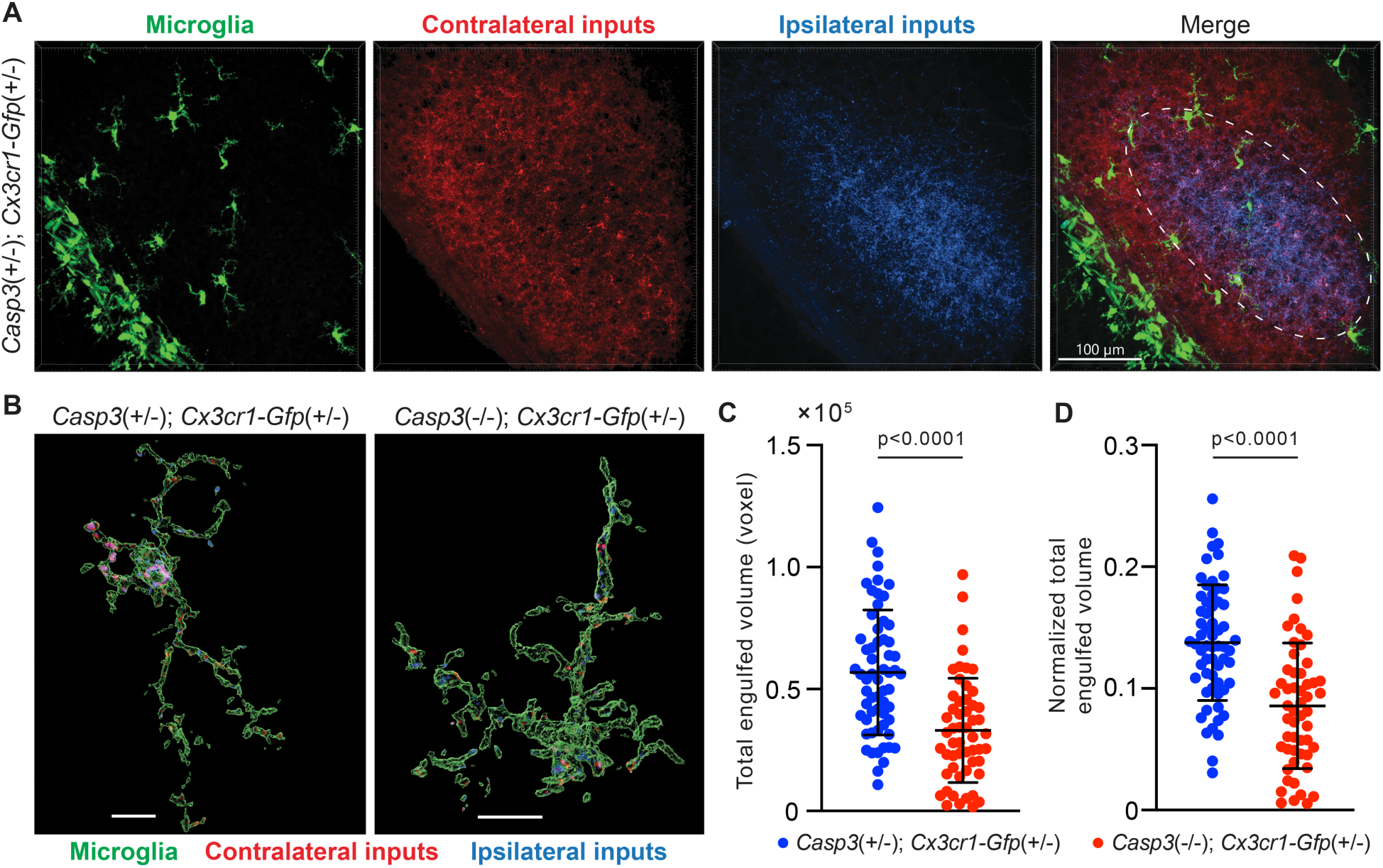
Microglia-mediated synapse elimination depends on caspase-3. (**A**) Representative 3D-reconstructed images of a P5 *Casp3*^+/−^; *Cx3cr1-Gfp*^+/−^ mouse dLGN with microglia displayed in green, contralateral RGC axon terminals in red, and ipsilateral RGC terminals in blue. In the merged image, the region from which microglia are selected for analysis is indicated with the dashed line. The scale-bar represents 100 μm. (**B**) Representative surface rendering of microglia (green) from P5 dLGNs of *Casp3*^+/−^; *Cx3cr1-Gfp*^+/−^ and *Casp3*^−/−^; *Cx3cr1-Gfp*^+/−^ mice. Intracellular contralateral (red) and ipsilateral (blue) RGC axon terminals are shown. Microglia from caspase-3 deficient mice engulf visibly less synaptic material. Scale-bars represent 10 μm. (**C**) Total volume of engulfed synaptic material in individual microglia from *Casp3*^+/−^; *Cx3cr1-Gfp*^+/−^ and *Casp3*^−/−^; *Cx3cr1-Gfp*^+/−^ mice. (**D**) Total volume of engulfed synaptic material in each microglia (from **C**) is normalized to the volume of that microglia. In **C**-**D**, each point represents one microglia. Mean and S.D. are shown. p-values were calculated from unpaired two-tailed t-tests. n=61 microglia from 8 *Casp3*^+/−^; *Cx3cr1-Gfp*^+/−^ mice and n=54 microglia from 5 *Casp3*^−/−^; *Cx3cr1-Gfp*^+/−^ mice.

To test whether caspase-3 activation is also required for astrocyte-mediated synapse elimination, we used the *Aldh1l1-Gfp* transgenic mouse line, where *Gfp* expression specifically labels astrocytes (Fig. S12A) (*49*). We again labeled RGC axon terminals in *Casp3*^+/−^; *Aldh1l1-Gfp*^+/−^ and *Casp3*^−/−^; *Aldh1l1-Gfp*^+/−^ littermate mice with CTB conjugates (Fig. S12A). Astrocytes were present in the P5 dLGN at a much higher density than microglia and possess very fine processes (Fig. S12A). To ensure segmentation fidelity, we only analyzed astrocyte cell bodies and their proximal processes (Fig. S12B). Unexpectedly, we found that the volume of axonal material engulfed by astrocytes was comparable in the two groups of mice (Fig. S12C). The same trend was observed when we normalized the volume of engulfed synaptic material to astrocyte volume (Fig. S12D-E). It is possible that, by using *Casp3*^+/−^ mice instead of *Casp3*^+/+^ mice as controls, we have obscured some defects in astrocyte-mediated synapse elimination. Nevertheless, given that astrocyte-mediated synapse elimination did not show clear dependence on caspase-3, we focused on measuring synapse engulfment by microglia as a readout of preferential elimination of weak synapses.

### Removal of weak synapses by microglia requires caspase-3 activity

If caspase-3 activation at weak synapses is important for synapse removal, then in caspase-3 deficient mice, microglia should lose the ability to engulf weak synapses. To test this hypothesis, we inactivated retinogeniculate synapses originating from right eyes in *Casp3*^+/+^ and *Casp3*^−/−^ mice with AAV-hSyn-TeTxLC injections (Fig. 6A). A group of animals was included for each genotype that received control injections. To measure synapse engulfment by microglia, we labeled RGC axon terminals with CTB conjugates and visualized microglia by immunostaining against Iba1, a marker specific for microglia in the brain (Fig. 6B-G). According to our model, in TeTxLC-injected *Casp3*^+/+^ mice, microglia should preferentially engulf inactive retinogeniculate synapses originating from the right eye (Fig. 6A), whereas in TeTxLC-injected *Casp3*^−/−^ mice, synapses from the right and the left eye should be engulfed at similar levels regardless of their strength (Fig. 6A).

**Figure 6.**
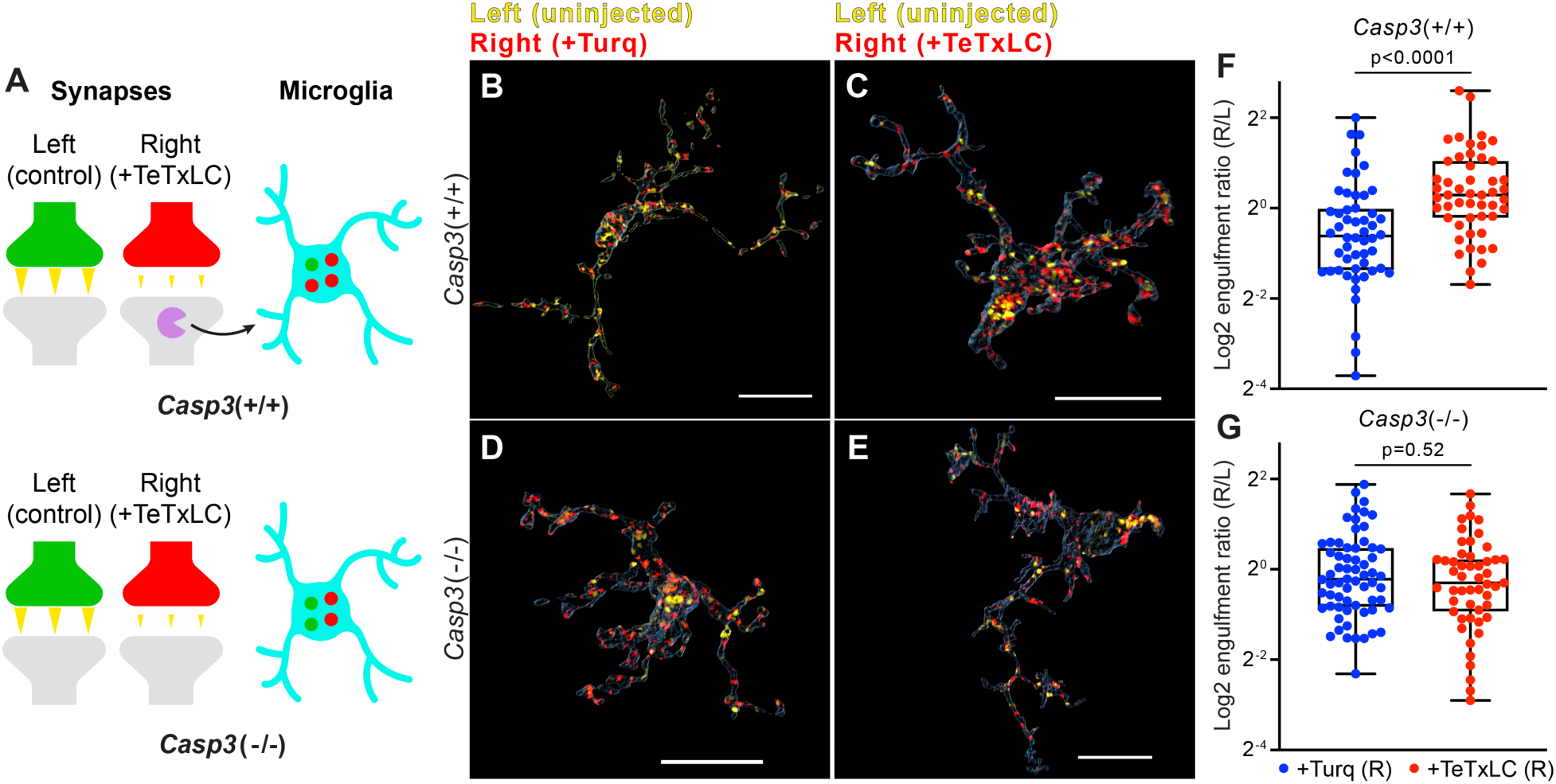
Removal of weak synapses by microglia requires caspase-3 activity. (**A**) Schematics illustrating the experimental rationale. In wildtype mice (upper panel), inactivating retinogeniculate synapses from the right eye activates caspase-3 (magenta) and recruits microglia to preferentially engulf right eye-originated synapses (red) over left eye-originated synapses (green). If caspase-3 activation is blocked (lower panel), engulfment of inactive synapses should be attenuated. (**B**-**E**) Surface rendering of representative microglia from P5 left dLGN of *Casp3*^+/+^ (**B**-**C**) or *Casp3*^−/−^ (**D**-**E**) mice injected with AAV that provided mTurquoise2 (**B** and **D**) or TeTxLC (**C** and **E**) in the right eye at E15. Microglia were labeled by immunostaining against Iba1. RGC axon terminals from the left eye are shown in yellow and terminals from the right eye in red. Scale bars represent 15 μm. (**F**-**G**) Ratio between volumes of right-eye and left-eye-originated synaptic material engulfed by microglia from *Casp3*^+/+^ (**F**) or *Casp3*^−/−^ (**G**) mice injected with AAV that carried the gene for mTurquoise2 (blue) or TeTxLC (red). Each dot represents one microglia. Engulfment ratios are displayed on a log scale. 0, 25, 50, 75, and 100 percentiles are shown. p-values were calculated from unpaired two-tailed Mann-Whitney tests. n=52 microglia from 7 Turq-injected *Casp3*^+/+^ mice, n=50 microglia from 8 TeTxLC-injected *Casp3*^+/+^ mice, n=64 microglia from 5 Turq-injected *Casp3*^−/−^ mice, and n=51 microglia from 6 TeTxLC-injected *Casp3*^−/−^ mice.

To quantitatively detect engulfment of weak synapses by microglia, we compared TeTxLC-injected and control-injected animals of the same genotype (Fig. 6B, C, and F). We first calculated the ratio between the volume of right eye-originated RGC axon terminals and left eye-originated terminals that were engulfed by each microglia. Then, by comparing this right-to-left engulfment ratio in TeTxLC-injected vs. control-injected animals, engulfment of inactive synapses could be detected as higher engulfment ratios in TeTxLC-injected animals (Fig. 6F). It is worth noting confounding factors, such as surgical manipulations, cancel out when calculating engulfment ratios, allowing information pertaining only to substrate preference to be isolated.

Consistent with previous studies (*7*), in *Casp3*^+/+^ mice inactivation of retinogeniculate synapses from the right eye led to enhanced microglia-mediated engulfment of right eye-originated RGC axon terminals but not left eye-originated terminals (Fig. S13A-B), resulting in a significant increase in right-to-left engulfment ratios in microglia of TeTxLC-injected animals compared to control-injected animals (Fig. 6B, C, and F). In contrast, engulfment ratios were similar between TeTxLC-injected and control-injected *Casp3*^−/−^ mice (Fig. 6D, E and G and Fig. S13C-D), suggesting that microglia could no longer distinguish between strong and weak synapses in the absence of caspase-3 activation. These observations remain valid when data were averaged and reported by animal (Fig. S13E-F). Summing the evidence obtained thus far, we propose the activity-dependent caspase-3 activation is a crucial signaling event that integrates upstream information about synaptic activity, takes place at weak/inactive synapses, and leads to activity-dependent elimination of weak synapses by microglia.

### Caspase-3 deficiency protects against amyloid-β-induced synapse loss

Lastly, we wanted to test if caspase-3 regulates synapse loss in neurodegenerative diseases through mechanisms analogous to its role in activity-dependent synapse elimination. To this end, we focused on Alzheimer’s disease (AD). AD is the most common cause of dementia (*50*), and synapse loss is the strongest correlate of cognitive decline in AD patients (*51*). Accumulation and deposition of oligomeric or fibril amyloid-β (Aβ) peptide has been proposed to be the cause of neurodegeneration in AD (*50*). Intriguingly, oligomeric Aβ impairs long-term potentiation (LTP) (*52*), and this suppression of LTP requires caspase-3 activation (*53*). Additionally, caspase-3 activation has been linked to early synaptic dysfunction in an AD mouse model (*54*).

To test if synapse loss in AD is regulated by caspase-3, we utilized the APP/PS1 mouse line, in which amyloid precursor protein (APP) and presenilin 1 (PS1) carrying mutations associated with early-onset familial AD are overexpressed in CNS neurons (*55*). We observed that female *App/Ps1*^+/−^ mice developed amyloid deposits in the hippocampus and cortex by 5 months (Fig. S14Aii), and that the amyloid burden worsened by 6 months (Fig. S14Aiii). In male *App/Ps1*^+/−^ mice, amyloid deposits only became apparent at 6 months and occurred at a lower level compared to age-matched females (Fig. S14Aiv-v). We focused on analyzing *App/Ps1*^+/−^ female mice in subsequent experiments to obtain robust phenotypes. To quantify synapse loss, we labeled presynaptic and postsynaptic compartments in the dentate gyrus of 6 month-old *App/Ps1*^−/−^ and *App/Ps1*^+/−^ littermates by detecting synaptic vesicle protein 2 (SV2) and Homer1, respectively, and selected multiple fields of interest per animal for imaging analysis while avoiding regions adjacent to amyloid deposits (Fig. S15). We fitted ellipsoids to SV2 and Homer1 signals to identify pre- and post-synaptic puncta and defined synapses as Homer1-SV2 puncta pairs that were less than 300 nm apart (Fig. 7A). We observed that synapse density in *App/Ps1*^+/−^ mice was significantly reduced compared to that in *App/Ps1*^−/−^ littermate controls (Fig. 7A-B), and that this reduction could be attributed to lower densities of both pre- and post-synaptic puncta in *App/Ps1*^+/−^ mice (Fig. S16A-B). We then crossed APP/PS1 mice to caspase-3 deficient mice and quantified synapse density in 6 month-old *Casp3*^−/−^; *App/Ps1*^−/−^ and *Casp3*^−/−^; *App/Ps1*^+/−^ littermate mice (Fig. S15). We observed that, in the caspase-3 deficient background, overexpression of mutant APP and PS1 no longer resulted in a reduction in the density of synapses (Fig. 7C-D) or pre-/post-synaptic puncta (Fig. S16C-D), suggesting that caspase-3 activity is required for Aβ-induced synapse loss in the APP/PS1 mouse model. Intriguingly, levels of amyloid deposition were comparable between *App/Ps1*^+/−^ and *Casp3*^−/−^; *App/Ps1*^+/−^ mice (Fig. S14Avi and B), and reactive microglia were observed near amyloid deposits regardless of caspase-3 deficiency (Fig. S17), indicating that protection conferred by caspase-3 deficiency against Aβ-induced synapse loss might not require inhibition of amyloid accumulation or microgliosis. We also investigated whether Aβ accumulation induced elevated levels of caspase-3 activity. We analyzed 4, 5, and 6 month-old *App/Ps1*^−/−^ and *App/Ps1*^+/−^ mice but only observed robust upregulation of caspase-3 activity in the dentate gyrus of a subset of 4 month-old *App/Ps1*^+/−^ mice (Fig. S18A-C). Upregulated caspase-3 activity in these mice remained in a punctate pattern without inducing neuronal death (Fig. S18D-F). It is possible that Aβ transiently induces caspase-3 activation in *App/Ps1*^+/−^ mice, but stochasticity in the onset and duration of such induction prevents reliable detection with our sample size and temporal resolution. Overall, our results demonstrate caspase-3 as an important regulator of Aβ-induced synapse loss and a potential therapeutic target for AD treatment.

**Figure 7.**
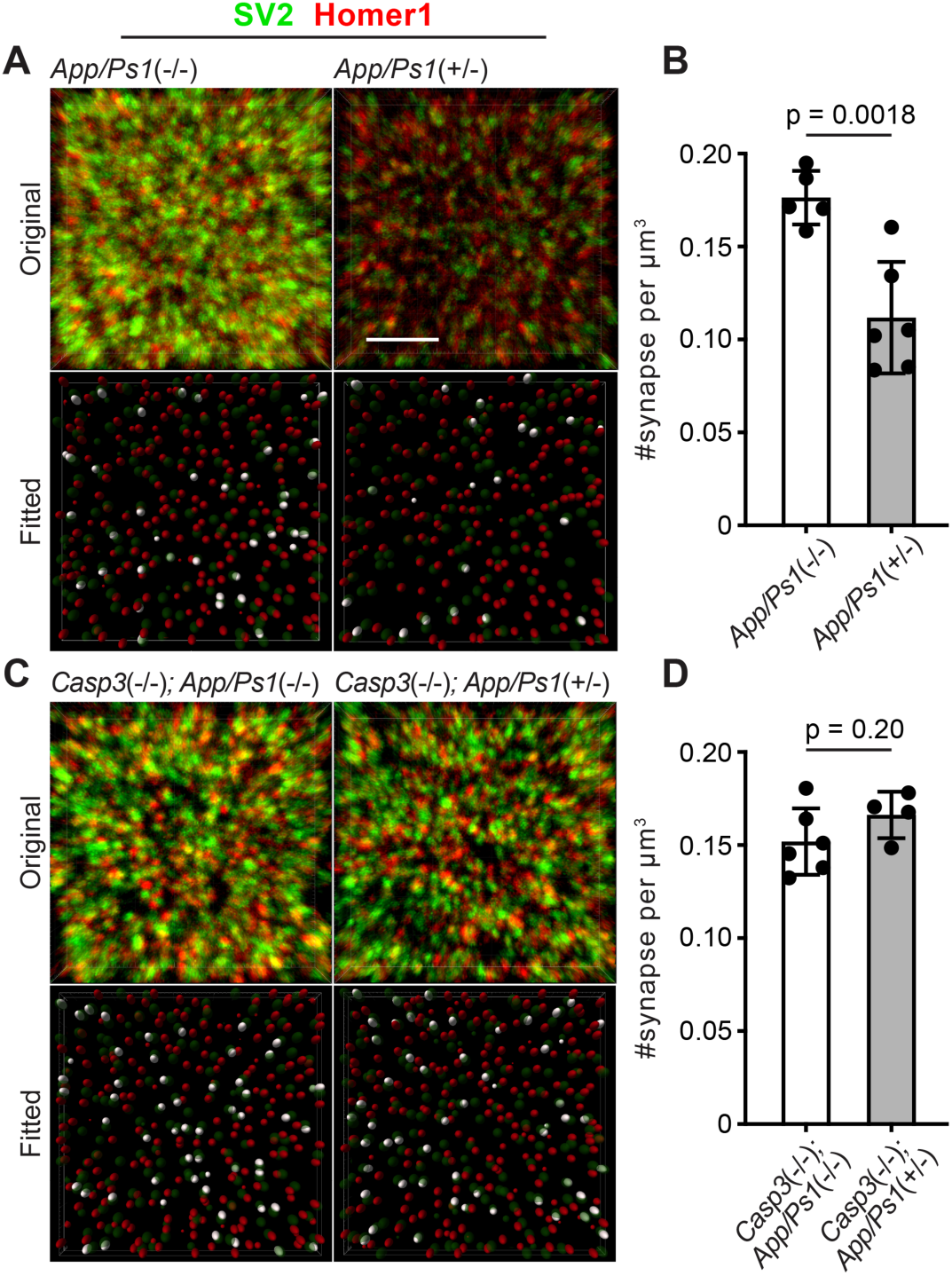
Caspase-3 deficiency protects against Aβ induced synapse loss. (**A** and **C**) Representative 3D-reconstructed images (3 μm in z) showing presynaptic (SV2, in green) and postsynaptic (Homer1, in red) signals in dentate gyrus of female 6 month-old APP/PS1 mice on caspase-3 wildtype (**A**) and deficient (**C**) backgrounds. Ellipsoids were fitted to the original images (upper panels) to isolate pre-(green) and post-synaptic (red) puncta (lower panels). Homer1 ellipsoids found with 300 nm of a SV2 ellipsoid are highlighted in white in the fitted images. Original images are adjusted to the same contrast. For the fitted images, only ellipsoids from the upper half of the z-stack are shown. Scale-bar represents 4 μm. (**B** and **D**) Quantification of synapse density in APP/PS1 mice on caspase-3 wildtype (**B**) and deficient (**D**) backgrounds. Mean and S.D. are shown. p-values were calculated from unpaired two-tailed t-tests. n=5 for *App/Ps1*^−/−^ mice, n=6 for *App/Ps1*^+/−^ mice, n=6 for *Casp3*^−/−^; *App/Ps1*^−/−^ mice, and n=4 for *Casp3*^−/−^; *App/Ps1*^+/−^ mice.

## Discussion

### A model for caspase-3 dependent synapse elimination

Our findings support a model where competitive interaction between strong synapses and weak synapses in the retinogeniculate pathway triggers caspase-3 activation in the postsynaptic compartments of weak synapses. Caspase-3 activation results in the removal of weaker synaptic connections by microglia, thereby facilitating the establishment of mature circuits in the visual pathway.

### Mechanisms of activity-dependent caspase-3 activation

In the experiment where we demonstrated activity-dependent caspase-3 activation, we inactivated all retinogeniculate synapses from one eye to induce sufficiently high caspase-3 activity that can be discerned from background. Widespread caspase-3 activity induced by such strong inactivation at multiple sites in a given cell likely self-amplifies through positive feedback (*35*) and overwhelms negative regulation mechanisms (e.g., ubiquitin-dependent proteasomal degradation (*24, 33, 34*)), leading to the observed apoptosis of entire dLGN relay neurons (Fig. 1B). However, apoptotic neurons were infrequently observed in the dLGN of control animals (Fig. 1D and 2B), and caspase-3 deficiency does not significantly alter dLGN relay neuron numbers (Fig. S7), suggesting that during normal development, activity-dependent caspase-3 activation is predominantly localized and generally does not lead to neuronal death.

Intriguingly, synaptic inactivation in our experiment was achieved through TeTxLC expression in presynaptic RGCs, but caspase-3 activation was observed in postsynaptic relay neurons. Conversely, while caspase-3 is activated in the postsynaptic compartment, we were able to detect defects in synapse elimination using assays that measure presynaptic inputs. These observations suggest that activity-dependent caspase-3 activation requires synaptic transmission, and that caspase-3 dependent synapse elimination is not strictly axonal or dendritic but involves both pre- and post-synaptic compartments. An important limitation of this study is that experiments in full-body *Casp3*^−/−^ mice cannot query the roles of caspase-3 in presynaptic and postsynaptic compartments separately. Future experiments that perturb caspase-3 activation and other signaling events specifically in the pre- or post-synaptic compartments are needed to reveal more detailed mechanisms of activity-dependent synapse elimination.

The observation that activity-dependent caspase-3 activation requires the presence of both weak and strong synapses resonates with previous research demonstrating that relative synaptic efficacy biases the outcome of synapse elimination during development of neuromuscular junctions and the callosal pathway (*15, 36, 56, 57*). However, the mechanisms linking synaptic competition with caspase-3 activation remains elusive. During LTD, caspase-3 is activated through mitochondrial release of cytochrome-c (*18*). Whether synaptic competition could induce caspase-3 activation via mechanisms such as calcium signaling, which regulates both neurotransmission and cytochrome-c release (*58*), is an outstanding question.

### Link between caspase-3 activation and synapse engulfment by microglia

How does caspase-3 activation at weak synapses lead to synapse engulfment by microglia? Caspase-3 activation is known to trigger the display of apoptosis markers that can be recognized by phagocytes (*19*). One of the best understood markers is phosphatidylserine (PtdSer), a phospholipid that translocates from the cytoplasmic leaflet of the plasma membrane to the exoplasmic leaflet during apoptosis (*19*). Importantly, caspase-3 directly regulates the enzymes that translocate PtdSer through enzymatic cleavage (*59, 60*). Two recent studies demonstrated that PtdSer is displayed on retinogeniculate synapses during postnatal development, and microglia use the exposed PtdSer as a cue to target and engulf synapses (*8, 9*). Whether PtdSer exposure is upregulated in response to synapse weakening and whether caspase-3 activity is required for such exposure are important questions that require further investigation.

The complement protein C1q has been observed to bind to PtdSer on apoptotic cells and facilitate phagocytic clearance (*19*). Additionally, a previous study found caspase-3 activity in C1q-positive synaptosomes (*61*), suggesting that C1q may bridge caspase-3 activation and synapse elimination. However, we investigated C1q expression and localization in the dLGN of P5 TeTxLC-injected mice but did not observe upregulation of C1q abundance or colocalization between C1q and caspase-3 activity (data not shown). It is often assumed in the literature that microglia directly engulf intact synapses, but existing data also support an alternative model where microglia indirectly contribute to synapse elimination by engulfing synaptic debris that is shed through neuron-autonomous mechanisms (*62*). The observation that synaptic activity promotes microglial engulfment through caspase-3 activation raises the possibility that weakened synapses partially disintegrate through apoptosis-like mechanisms prior to being engulfed by microglia. Further studies are needed to elucidate the intermediate steps between caspase-3 activation and microglial engulfment of synapses.

### Caspase-3 as a potential target for AD treatment

Our findings suggest that caspase-3 deficiency confers protection against Aβ-induced synapse loss in a mouse model of AD. Notably, the absence of caspase-3 does not seem to affect Aβ deposition or microglia reactivity, indicating that caspase-3 depletion may mitigate synapse loss by directly countering Aβ-induced synaptic toxicity. Given the prominence of Aβ and inflammation as targets for AD drug development (*63*), inhibiting caspase-3 activity offers a potential alternative therapeutic strategy that could complement and enhance current treatment approaches. Therefore, it is important to test whether pharmacological inhibition of caspase-3 can effectively slow synapse loss and cognitive decline in both mouse models of AD and human patients.

## Acknowledgement

We thank Crystall Lopez, Benjamin Gantz, and Brooke Groff for help with animal husbandry and breeding. We thank Sarah Lindo, Colin Morrow, Claire Boyer, and Rae Demars for help with surgery. We thank Monique Copeland, Amy Hu, Morgan Clarke, and Benjamin Foster for help with histology. We are also grateful to Sarah Kivimaki and the Viral Tools team of Janelia Core Services for help with AAV preparation. We thank Dr. Cagla Eroglu, Dr. Richard Axel, and Dr. David Clapham for comments on the manuscript. This work is supported by the Howard Hughes Medical Institute.

