## supplementary figures for "Activity-dependent synapse elimination requires caspase-3 activation"

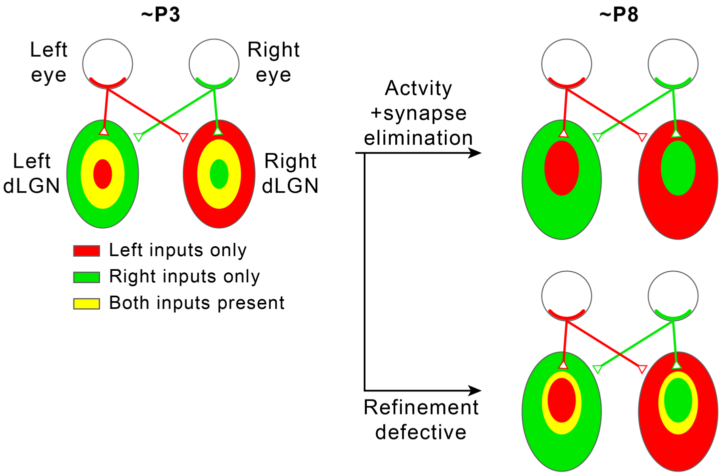


**Figure S1. Segregation of eye-specific territories in the mouse retinogeniculate pathway.** In the mouse retinogeniculate pathway, retinal ganglion cells (RGCs) in the retina of each eye innervate relay neurons in both the contralateral (opposite side as the originating RGC) dorsal lateral geniculate nucleus (dLGN) and the ipsilateral (same side from the originating RGC) dLGN to form retinogeniculate synapses (upper right). Within each dLGN, the majority of retinogeniculate synapses receive inputs from the contralateral eye, while the minority receive inputs from the ipsilateral eye (upper right). At the age of P3, regions in each dLGN receiving inputs from each of the two eyes overlap significantly (left). Through a process that requires synapse elimination and spontaneous RGC activity, these regions are refined into non-overlapping eye-specific territories by the age of P8 (upper right). If the refinement process is defective because neural activity or synapse elimination is disrupted, eye-specific territories fail to completely segregate, and regions innervated by the two eyes remain overlapping (lower right).

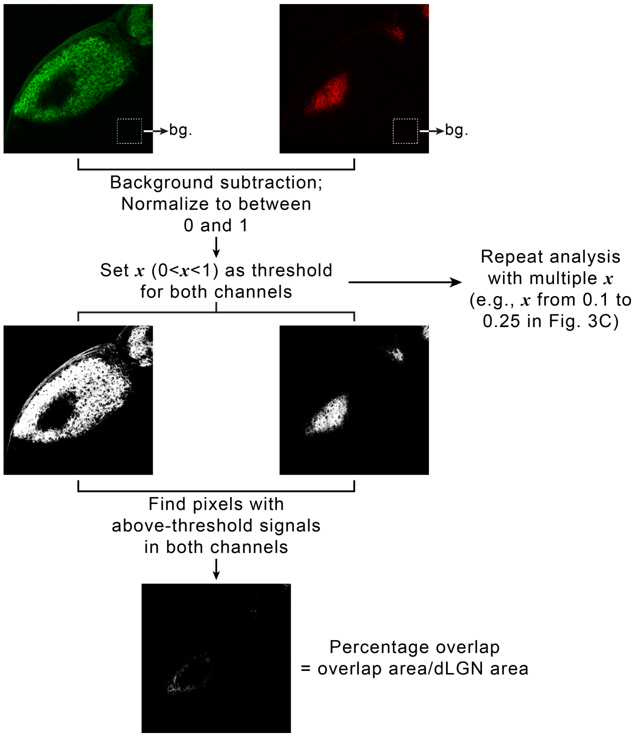


**Figure S2. Quantifying eye-specific segregation with multi-threshold overlap analysis.** For each dLGN, RGC inputs from the two eyes were imaged using separate fluorescence channels. A small area in the thalamus outside of each dLGN was chosen, and average signal intensity in each channel within that area was calculated and used as background (upper panel). For each channel, background was subtracted, and signals were normalized to between 0 and 1. To calculate overlap between eye-specific territories, a threshold, $x$, was chosen between 0 and 1 and applied to both channels (middle panel). The overlap between eye-specific territories were defined as the set of pixels with above-threshold signals in both channels. Percentage overlap was then calculated as the ratio between the area of the dLGN where eye-specific territories overlapped and the total area of the dLGN (lower panel). To avoid introducing biases by artificially selecting one threshold, we repeated the analysis with a set of increasingly stringent thresholds (e.g., from 0.1 to 0.25 in Fig. 3C and 3D).

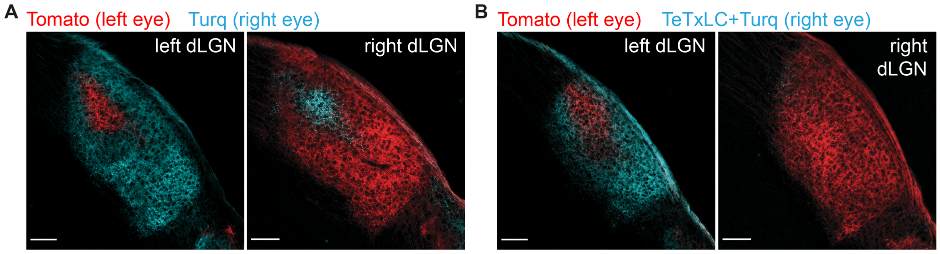


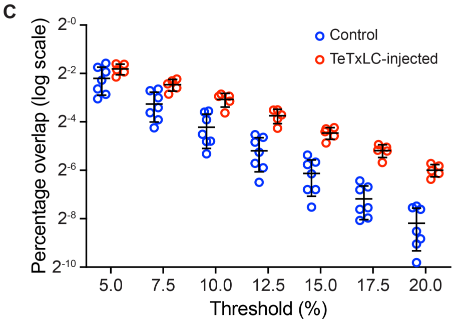


**D**

| Threshold (%) | Control mean | TeTxLC mean | P-value |
|---|---|---|---|
| 5.0 | 0.2172 | 0.2839 | 0.135 |
| 7.5 | 0.1045 | 0.1809 | 0.00601 |
| 10.0 | 0.05357 | 0.1187 | 0.00089 |
| 12.5 | 0.02736 | 0.07447 | 0.00014 |
| 15.0 | 0.01427 | 0.04548 | 0.000022 |
| 17.5 | 0.006909 | 0.02745 | 0.000005 |
| 20.0 | 0.003429 | 0.01568 | 0.000003 |

**Figure S3. Mono-ocular blockage of RGC activity with TeTxLC disrupts eye-specific segregation.** (**A** and **B**) Confocal images of P8 dLGNs in animals receiving either control injections (**A**) or TeTxLC injections in the right eye (**B**). RGC inputs were labeled with tdTomato (red, input from left eyes) or mTurquiose2 (cyan, input from right eyes) to visualize eye-specific territories. The territory of right eye inputs (cyan) contracted relative to that of left eye inputs (red) in TeTxLC-injected animals. Scale-bars: 100 μm. (**C**) Quantification of overlap between eye-specific territories in dLGNs of control and TeTxLC-injected animals. Analysis was done at multiple thresholds to avoid biases introduced with threshold selection. Percentage overlap is displayed on a log scale. n=7 for control animals and 5 for TeTxLC-injected animals. Mean and S.D. are shown. (**D**) Statistics of the multi-threshold analysis showing the percentage overlap in the two groups of animals at each threshold. P-values for the difference between control and TeTxLC-injected animals were calculated from two-tailed t-tests. Note that the statistical significance increased with more stringent thresholds. We did not implement multiple comparison corrections as values at different thresholds are derived from the same dataset and are not independent.

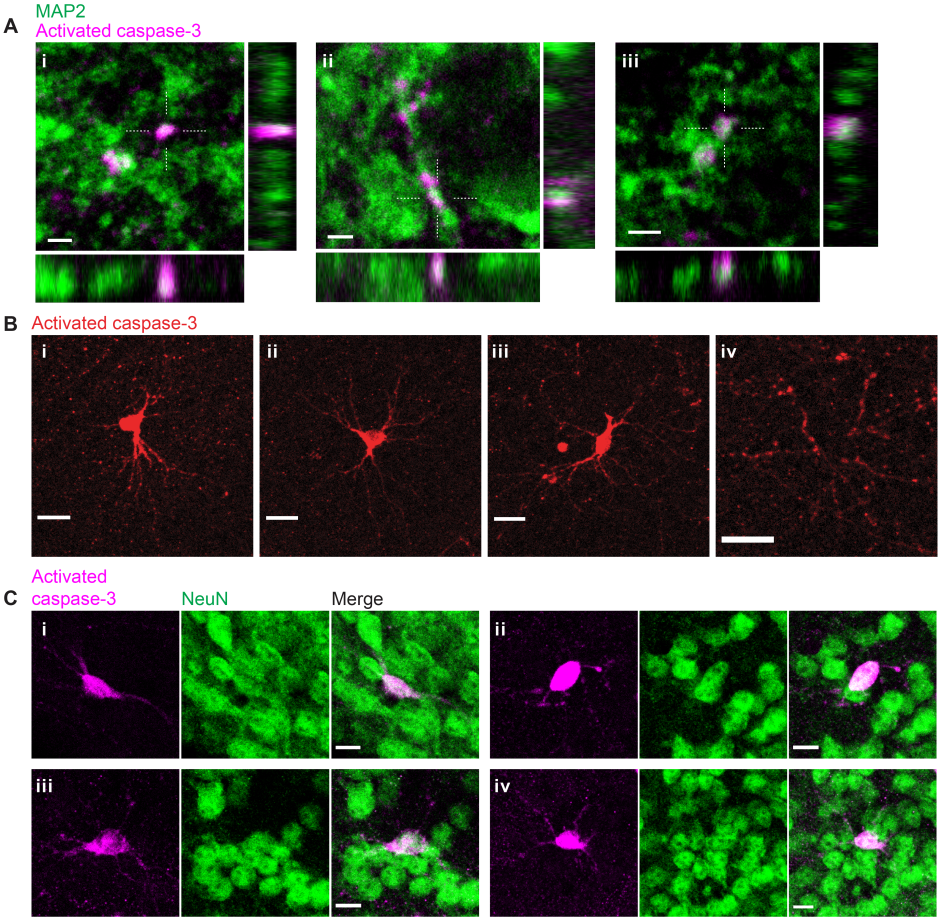


**Figure S4. Inactivation of retinogeniculate synapses induces postsynaptic caspase-3 activity in dendritic compartments of dLGN relay neurons.** (**A**) High-resolution images of three representative field-of-views (**i-iii**) in P5 TeTxLC-expressing dLGNs showing co-localization of punctate caspase-3 activity (magenta) and a dendritic marker, MAP2 (green). Dotted crosses mark x and y positions where the x-z and y-z cross-sections were generated. Scale-bars: 2 μm. (**B**) Example images showing caspase-3 activity in entire neurons (**i-iii**) or in multiple dendritic branches (but not in the soma) of a neuron (**iv**) in P5 TeTxLC-expressing dLGNs. The neurons positive for active caspase-3 (**i-iii**) have relatively large and round somas and multipolar dendritic arbors that are characteristic of dLGN relay neurons. Scale-bars: 20 μm. (**C**) Example images of four representative neurons (**i-iv**) in P5 TeTxLC-expressing dLGNs that are positive for active caspase-3 (magenta) and a neuronal nuclear marker, NeuN (green). Scale-bars: 10 μm.

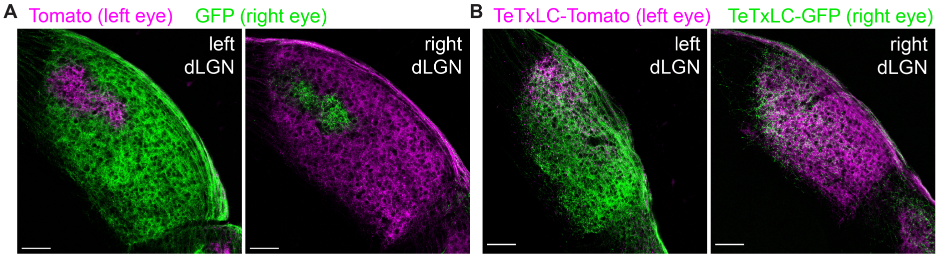


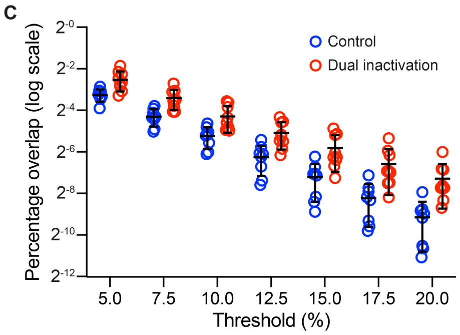


**D**

| Threshold (%) | Control mean | Dual inact. mean | P-value |
|---|---|---|---|
| 5.0 | 0.1043 | 0.1732 | 0.005 |
| 7.5 | 0.0508 | 0.0940 | 0.0024 |
| 10.0 | 0.0265 | 0.0512 | 0.009 |
| 12.5 | 0.0131 | 0.0294 | 0.0044 |
| 15.0 | 0.0067 | 0.0177 | 0.0086 |
| 17.5 | 0.0033 | 0.0104 | 0.012 |
| 20.0 | 0.0018 | 0.0064 | 0.0072 |

**Figure S5. Bi-ocular blockage of RGC activity with TeTxLC disrupts eye-specific segregation.** (**A** and **B**) Confocal images of P10 dLGNs of animals receiving control injections in both eyes (**A**) or TeTxLC injections in both eyes (**B**). Eye-specific territories were labeled with tdTomato (magenta, RGC inputs from the left eye) or eGFP (green, RGC inputs from the right eye). Scale-bars: 100 μm. (**C**) Quantification of overlap between eye-specific territories in dLGNs of animals receiving control or TeTxLC injections in both eyes. n=8 for control animals and n=9 for dually TeTxLC-injected animals. Mean and S.D. are shown. (**D**) Statistics of the overlap analysis at multiple thresholds. P-values were calculated from two-tailed t-tests. We did not implement multiple comparison corrections as values at different thresholds are derived from the same dataset and are not independent.

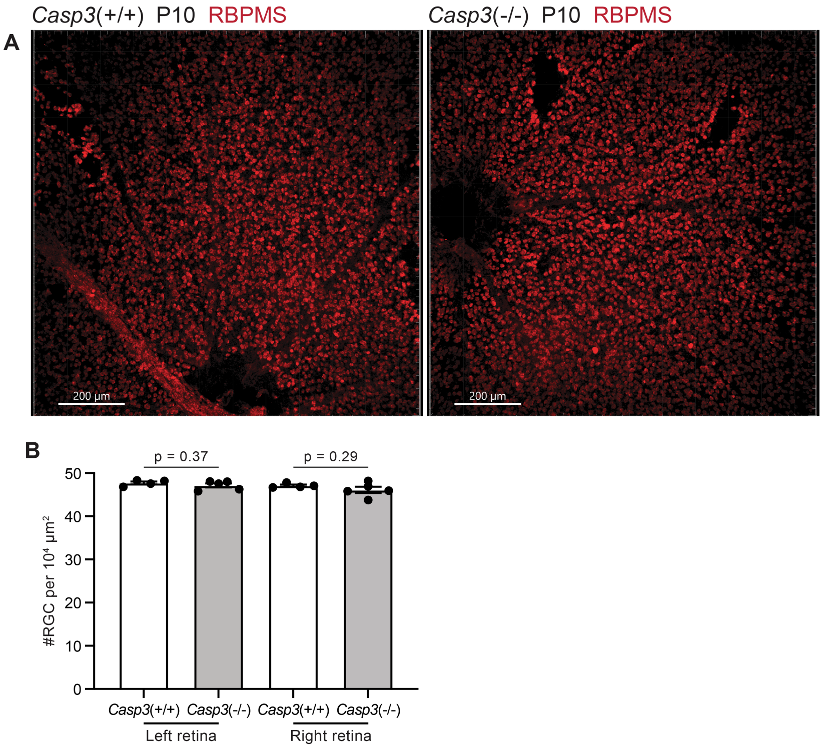


**Figure S6. Caspase-3 deficiency does not alter RGC density in the retina.** (**A**) Images of whole-mount retinae from P10 *Casp3*$^{+/+}$ (left) and *Casp3*$^{-/-}$ (right) animals. RGCs were labeled in red by immunostaining against an RGC-specific marker, RBPMS (RNA-binding protein with multiple splicing) (*37*). The dark regions on the bottom of the left panel and on the left of the right panel are optic discs. Scale-bars: 200 μm. (**B**) Quantification of RGC densities in retinae of P10 *Casp3*$^{+/+}$ and *Casp3*$^{-/-}$ animals. Each point represents one retina. Left and right retinae were analyzed separately. n=4 animals for *Casp3*$^{+/+}$ mice and n=5 animals for *Casp3*$^{-/-}$ mice. Mean and S.D. are shown. P-values were calculated from unpaired two-tailed t-tests.

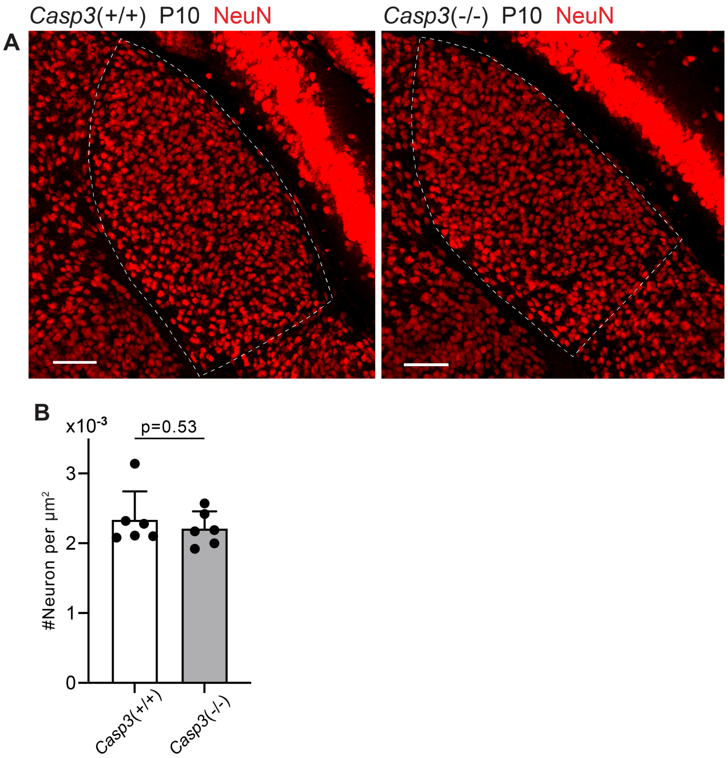


**Figure S7. Caspase-3 deficiency does not alter relay neuron density in the dLGN.** (**A**) Images of dLGNs of P10 *Casp3*$^{+/+}$ (left) and *Casp3*$^{-/-}$ (right) animals. Dashed lines mark the dLGN boundaries. Relay neurons were labeled by immunostaining against NeuN. Scale-bars: 100 μm. (**B**) Quantification of dLGN relay neuron densities in P10 *Casp3*$^{+/+}$ and *Casp3*$^{-/-}$ mice. Each point represents one animal. n=6 animals for *Casp3*$^{+/+}$ mice and n=6 animals for *Casp3*$^{-/-}$ mice. Mean and S.D. are shown. P-values were calculated from unpaired two-tailed t-tests.

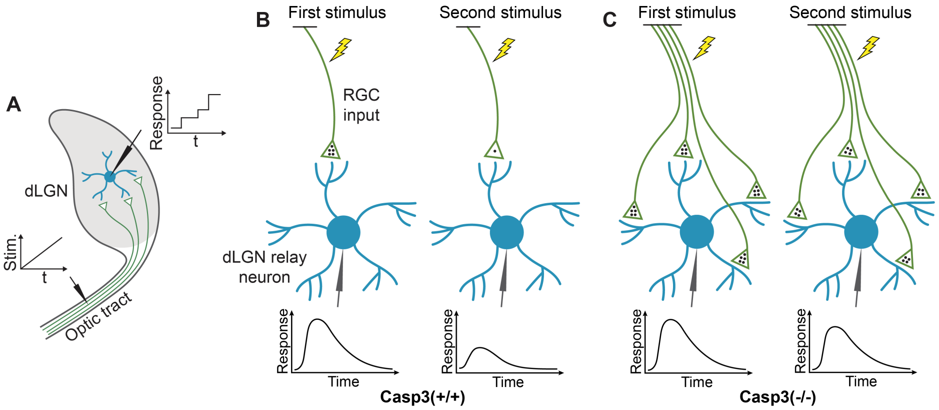


**Figure S8. Measuring electrophysiological properties of retinogeniculate synapses.** (**A**) Inferring the number of RGC inputs of dLGN relay neurons by measuring stimulation-response curves. To measure relay neuron responses, we prepared acute brain slices from a tilted parasagittal plane in P30 mice (*38*). These slice preparations preserved a long segment of the optic tract and a high level of connectivity between RGC axons and dLGN relay neurons (*38*). We placed a stimulating electrode on the optic tract and patch-clamped individual dLGN relay neurons with a recording electrode. We stimulated the optic tract with a series of gradually increasing currents and recorded excitatory postsynaptic currents (EPSCs) in relay neurons. At low stimulation intensity, no RGC axons were activated. As the stimulus increased, more and more RGC axons were recruited. As different RGC axons have different excitation thresholds, RGC axons were excited one at a time. Whenever a stimulus activated an RGC axon that did not respond to lower stimuli, a step increase in the relay neuron response was detected, and the increment corresponded to the EPSC evoked by the newly recruited RGC axon. We can therefore infer the number of RGC inputs innervating the relay neuron being recorded by counting the number of steps in the response curve of that relay neuron. t: time. Stim.: stimulation intensity. (**B**-**C**) Inferring the number of release sites from RGC inputs by measuring paired-pulse response ratio (PPR). We stimulated the optic tract with two stimuli separated by a short interval and recorded EPSCs in dLGN relay neurons. The stimulus intensity was chosen to evoke maximum response in the relay neuron. In wildtype mice (**B**), the first stimulus triggers the release of a significant fraction of the readily releasable pool (RRP) of neurotransmitters (black dots), evoking a strong response (left). At the time of the second stimulus, the RRP does not have sufficient time to recover and remains depleted, resulting in a weaker second response (right). PPR can be calculated from this experiment by dividing the peak amplitude of the second response with that of the first. PPR is small in wildtype animals. In caspase-3 deficient mice (**C**), if the number of release sites is increased compared to wildtype animals, the RRP should be larger (left), and a smaller fraction of the RRP is released during the first stimulus, leaving more neurotransmitters available for the second stimulus (right), thereby enhancing the second response and PPR. Note that in this model, we made two assumptions. The first assumption is that the average size of the RRP in one release site is similar in *Casp3*$^{+/+}$ and *Casp3*$^{-/-}$ mice, which is supported by the comparable fiber fractions (FFs) in the two groups of animals (see Fig. S9E). The second assumption is that the number of neurotransmitters released during the first stimulus is similar in *Casp3*$^{+/+}$ and *Casp3*$^{-/-}$ mice (4 dots in the illustration), which is supported by the comparable maximum EPSCs in the two groups of animals (see Fig. S9B-C).

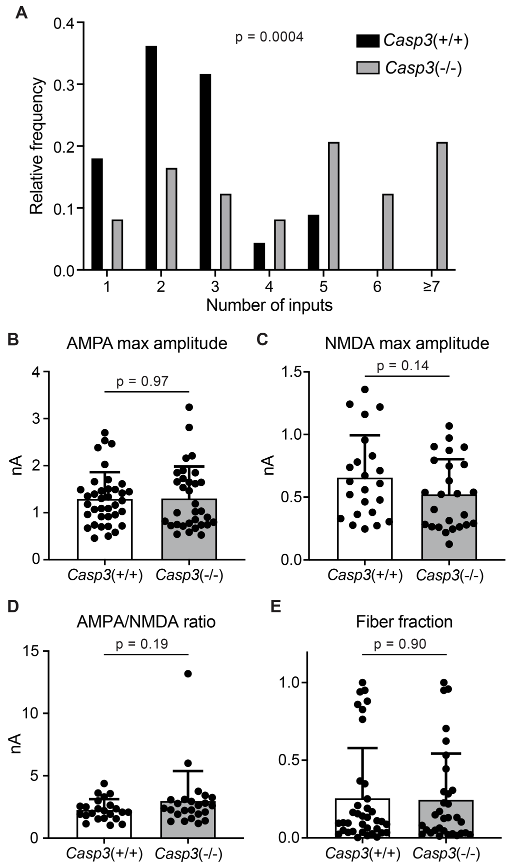


**Figure S9. Additional analyses of electrophysiological properties of retinogeniculate synapses in *Casp3*$^{+/+}$ and *Casp3*$^{-/-}$ mice.** (**A**) Distribution of RGC input numbers of individual dLGN relay neurons in wildtype and *Casp3*$^{-/-}$ mice measured by counting the number of steps in NMDAR-mediated EPSC response curves (upper right in Fig. 4A and 4B) while blind to genotypes. P-value was calculated from two-tailed t-test. n=22 cells for wildtype mice and n=24 cells for *Casp3*$^{-/-}$ mice. (**B**-**C**) Quantification of maximum amplitudes of AMPAR- (**B**) and NMDAR-mediated (**C**) EPSCs in wildtype and *Casp3*$^{-/-}$ dLGN relay neurons. (**D**) Quantification of the ratio between maximum amplitudes of AMPAR- and NDMAR-mediated EPSCs in wildtype and *Casp3*$^{-/-}$ dLGN relay neurons. (**E**) Quantification of fiber fractions in wildtype and *Casp3*$^{-/-}$ dLGN relay neurons. When stimulation on the optic tract is gradually increased, there will be a lowest stimulation intensity at which a non-zero response is first recorded in the relay neuron (see Fig. S8 for illustration and Fig. 4 for example data). This first response is presumed to be evoked by the activation of a single RGC axon fiber. Fiber fraction is defined as the ratio between this single-fiber response and the maximum response and ranges from 0 to 1. Only the AMPAR-mediated response is used to calculate fiber fractions. In **B**-**E**, mean and S.D. are shown. In **A**-**E**, p-values were calculated from two-tailed t-tests. In **A**, **C**, and **D**, n = 22 cells for wildtype mice and n=24 cells for *Casp3*$^{-/-}$ mice. In **B** and **E**, n=37 cells for wildtype mice and n=32 cells for *Casp3*$^{-/-}$ mice.

Deleted: ¶

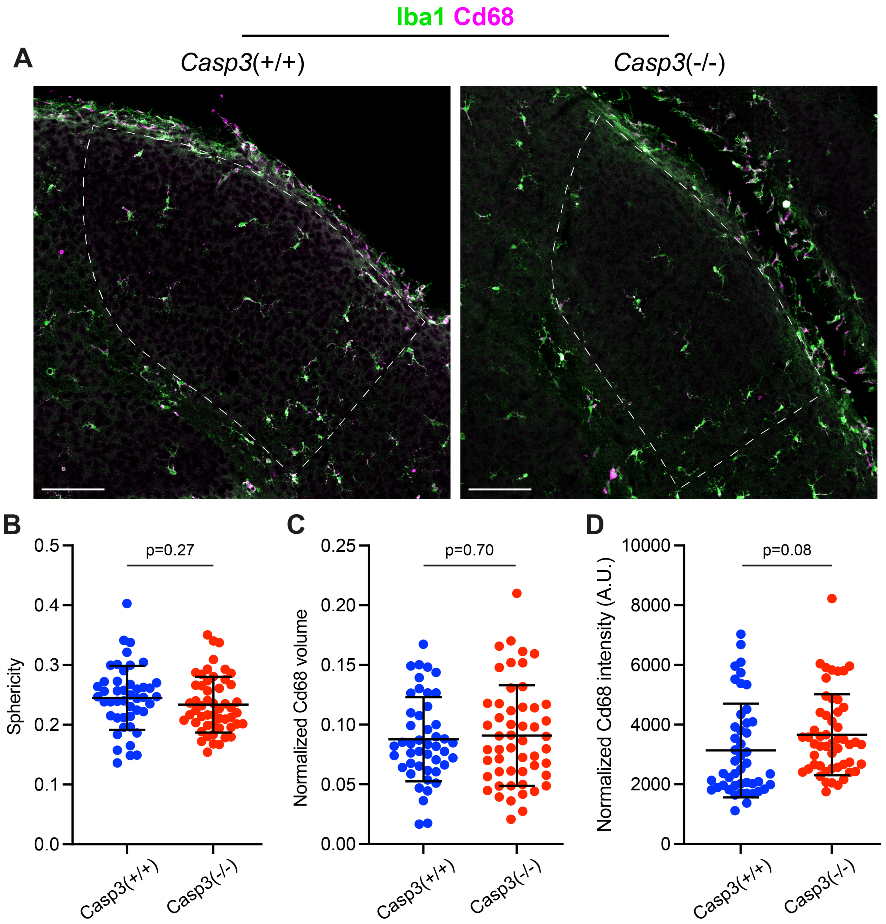


**Figure S10. Caspase-3 deficiency does not cause microglia activation.** (**A**) Representative images of P5 dLGNs in a *Casp3*$^{+/+}$ mouse (left) and a *Casp3*$^{-/-}$ mouse (right) showing Iba1 (green, a marker for microglia cell body) and Cd68 (magenta, a marker for microglia activation) staining. The dLGN areas are highlighted with dotted lines. Scale-bars represent 100 μm. (**B**) Quantification of microglia morphology in *Casp3*$^{+/+}$ and *Casp3*$^{-/-}$ mice using the sphericity metric. A sphericity of 1 corresponds to a perfect sphere. The smaller the sphericity is, the more ramified the cell is. (**C**-**D**) Quantification of microglia activation by normalizing the total volume of Cd68 signal (**C**) or the total intensity of Cd68 signal (**D**) in each microglia to the volume of that microglia. By both morphology and Cd68 signal, microglia in *Casp3*$^{-/-}$ mice show no evidence of activation. In **B**-**D**, each data point represents one microglia, and mean and S.D. are shown. n=46 microglia from 4 *Casp3*$^{+/+}$ mice and n=52 microglia from 4 *Casp3*$^{-/-}$ mice. P-values were calculated from two-tailed unpaired t-tests.

Deleted: ¶

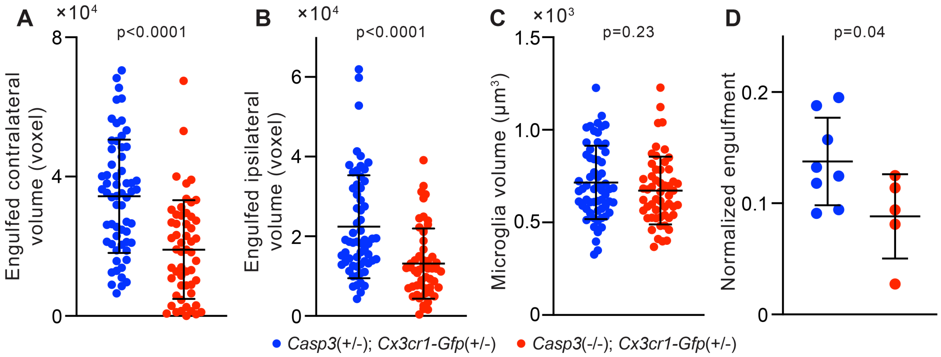


**Figure S11. Additional analyses of microglia-mediated engulfment of synaptic material.** (**A**-**B**) Volume of contralateral (**A**) and ipsilateral (**B**) RGC axon terminals engulfed by individual microglia in the P5 dLGN of *Casp3*$^{+/-}$; *Cx3cr1-Gfp*$^{+/-}$ and *Casp3*$^{-/-}$; *Cx3cr1-Gfp*$^{+/-}$ mice. (**C**) Volume of individual microglia in the same animals as in **A**-**B**. (**D**) Normalized engulfment values as in Figure 5D but averaged by animal. In **A**-**C**, each point represents one microglia. In **D**, normalized engulfment values for all microglia in each mouse were averaged and reported as one data point. Mean and S.D. are shown. P-values were calculated from unpaired two-tailed t-tests. n=61 microglia from 8 *Casp3*$^{+/-}$; *Cx3cr1-Gfp*$^{+/-}$ mice and n=54 microglia from 5 *Casp3*$^{-/-}$; *Cx3cr1-Gfp*$^{+/-}$ mice.

Deleted: ¶

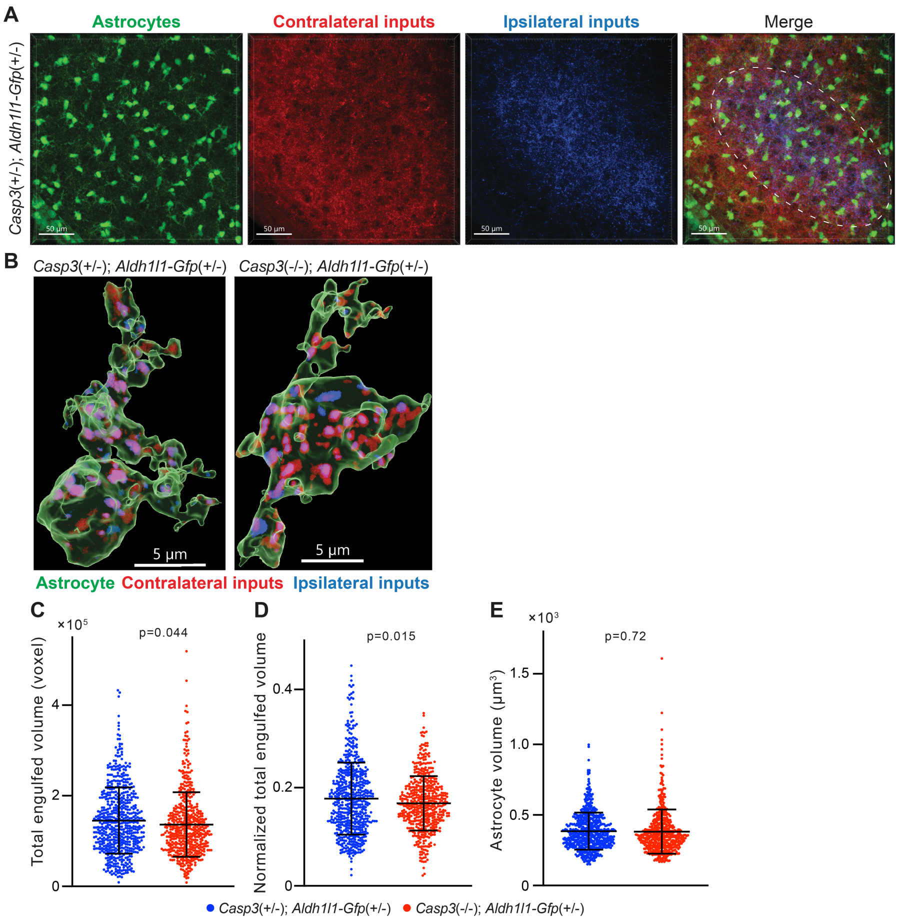


**Figure S12. Astrocyte-mediated synapse elimination does not appear to depend on caspase-3.** (**A**) Representative 3D-reconstructed images of a P5 *Casp3*$^{+/-}$; *Aldh1l1-Gfp*$^{+/-}$ mouse dLGN with astrocytes displayed in green, contralateral RGC axon terminals in red, and ipsilateral RGC terminals in blue. The region from which astrocytes are selected for analysis is indicated with the dashed line in the merged image. Scale-bars represent 50 µm. (**B**) Representative surface rendering of astrocytes (green) from P5 dLGNs of *Casp3*$^{+/-}$; *Aldh1l1-Gfp*$^{+/-}$ and *Casp3*$^{-/-}$; *Aldh1l1-Gfp*$^{+/-}$ mice. Intracellular contralateral (red) and ipsilateral (blue) RGC axon terminals are shown. Only cell bodies and the base of processes were segmented. We made this segmentation choice because we observed that fine astrocytic processes are largely devoid of engulfed material. Scale-bars represent 5 µm. (**C**) Total volume of engulfed synaptic material in individual astrocytes from *Casp3*$^{+/-}$; *Aldh1l1-Gfp*$^{+/-}$ and *Casp3*$^{-/-}$; *Aldh1l1-Gfp*$^{+/-}$ mice. (**D**) Total volume of engulfed synaptic material in each astrocyte (from **C**) is normalized to the volume of that astrocytes. (**E**) Volume of individual astrocytes from the two groups of mice. In **C**-**E**, each point represents one astrocyte. Mean and S.D. are shown. P-values were calculated from unpaired two-tailed t-tests. n=595 astrocytes from 8 *Casp3*$^{+/-}$; *Aldh1l1-Gfp*$^{+/-}$ mice and n=517 astrocytes from 6 *Casp3*$^{-/-}$; *Aldh1l1-Gfp*$^{+/-}$ mice.

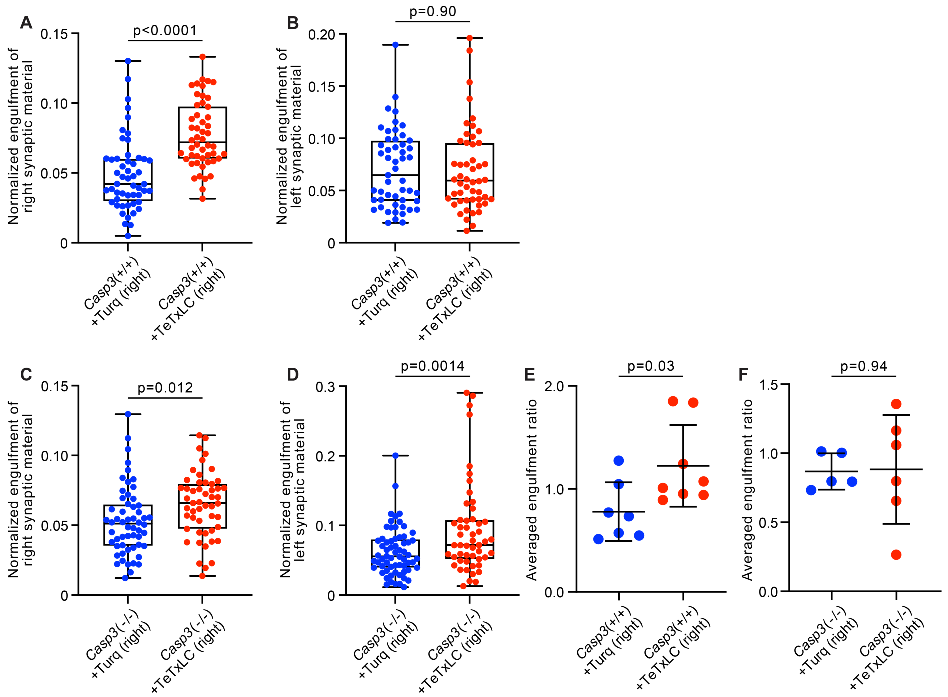


**Figure S13. Additional analysis on activity-dependent microglia-mediated engulfment of synapses.** (**A**-**D**) Volumes of synaptic material originating from right (**A** and **C**) or left (**B** and **D**) eyes engulfed by individual microglia from the left dLGN of *Casp3*$^{+/+}$ (**A**-**B**) or *Casp3*$^{-/-}$ (**C**-**D**) mice injected with AAV expressing mTurquoise2 (blue) or TeTxLC (red) in right eye were normalized to microglial volume and plotted. In *Casp3*$^{+/+}$ mice, inactivating right eye-originated retinogeniculate synapses specifically enhanced microglia-mediated engulfment of RGC axon terminals from right eyes (**A**) but not left eyes (**B**), resulting in significantly higher engulfment ratios (Fig. 6F). In *Casp3*$^{-/-}$ mice, inactivating right eye-originated synapses resulted in no significant change in engulfment ratios (Fig. 6G), even though engulfment of RGC terminals from both eyes was elevated (**C** and **D**). (**E**-**F**) Engulfment ratios as reported in Figure 6F-G but averaged by animal. In **A**-**D**, each point represents one microglia. 0, 25, 50, 75, and 100 percentiles are shown. In **E**-**F**, engulfment ratios for all microglia in each animal were averaged and reported as one data point. p-values were calculated from unpaired two-tailed t-tests. n=52 microglia from 7 Turq-injected *Casp3*$^{+/+}$ mice, n=50 microglia from 8 TeTxLC-injected *Casp3*$^{+/+}$ mice, n=64 microglia from 5 Turq-injected *Casp3*$^{-/-}$ mice, and n=51 microglia from 6 TeTxLC-injected *Casp3*$^{-/-}$ mice.

***Notes on interpretation of C and D*:** In *Casp3*$^{-/-}$ brains injected with TeTxLC, microglia in dLGNs tended to increase in volume and have thicker processes. We therefore suspect the non-specific upregulation of synapse engulfment in TeTxLC-injected *Casp3*$^{-/-}$ mice is a consequence of microglial activation rather than synapse inactivation. This microglial activation is likely a side effect of intraocular surgeries. We noticed that *Casp3*$^{-/-}$ brains tended to have weak structural integrity. It is possible that repetitive intraocular injections caused damage in the retinogeniculate pathway of *Casp3*$^{-/-}$ mice and activated microglia. Nevertheless, by calculating right-to-left engulfment ratios, substrate preference of microglia in TeTxLC-injected *Casp3*$^{-/-}$ mice can be deduced.

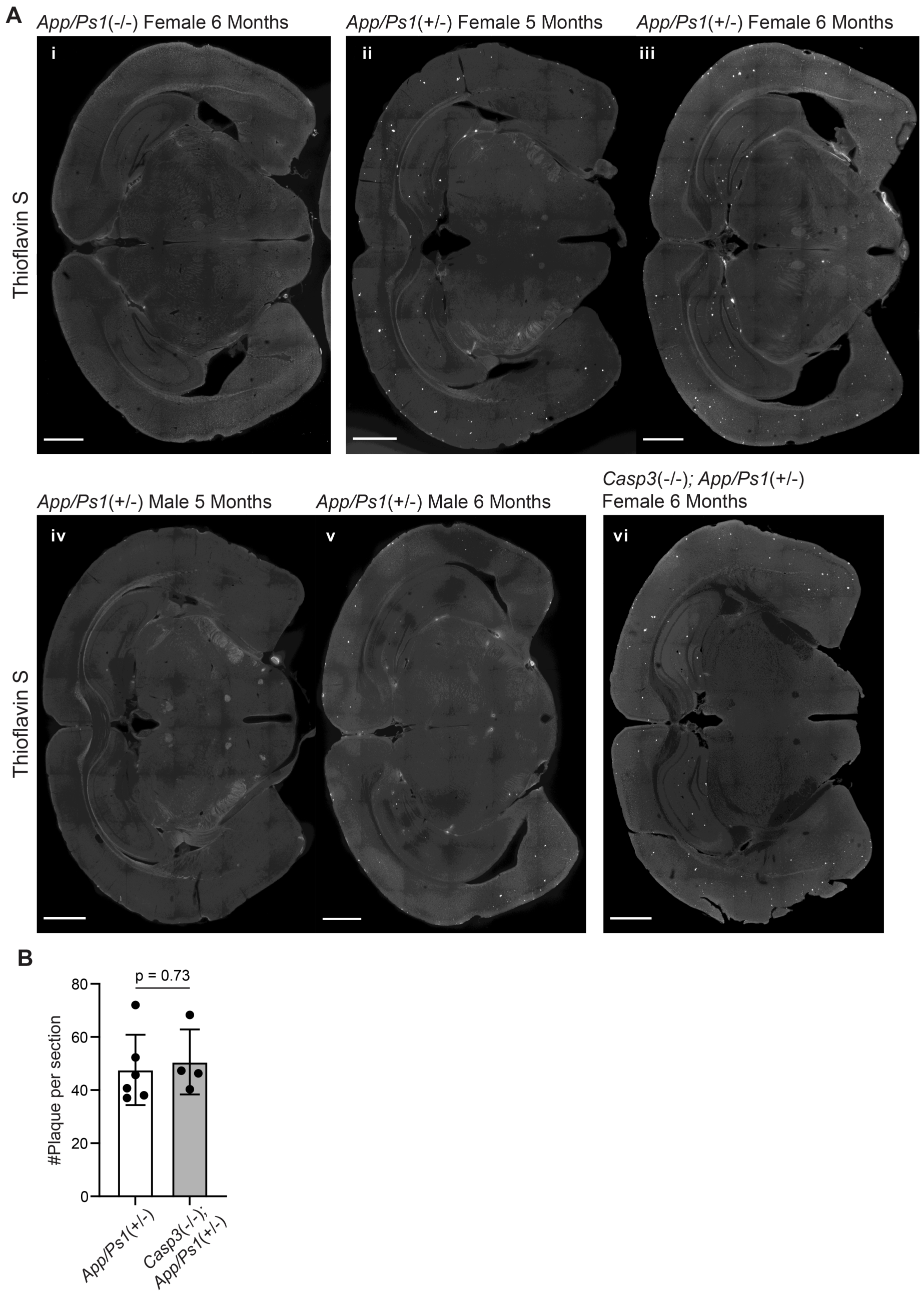
A
App/Ps1(-/-) Female 6 Months
App/Ps1(+/-) Female 5 Months
App/Ps1(+/-) Female 6 Months
i
ii
iii
Thioflavin S
App/Ps1(+/-) Male 5 Months
App/Ps1(+/-) Male 6 Months
Casp3(-/-); App/Ps1(+/-)
Female 6 Months
iv
v
vi
Thioflavin S
B
p = 0.73
#Plaque per section
80
60
40
20
0
App/Ps1(+/-)
Casp3(-/-);
App/Ps1(+/-)

**Figure S14. Amyloid deposition APP/PS1 mouse lines.** (**A**) Images of coronal brain sections of a female *App/Ps1*$^{-/-}$ mouse at 6 months (i), female *App/Ps1*$^{+/-}$ mice at 5 months (ii) or 6 months (iii), male *App/Ps1*$^{+/-}$ mice at 5 months (iv) or 6 months (v), and a female *Casp3*$^{-/-}$; *App/Ps1*$^{+/-}$ mouse at 6 months (vi), stained with Thioflavin S to reveal Aβ plaques (bright spots in hippocampus and cortex). Scale-bars represent 1 mm. (**B**) Quantification of the number of plaques per section in female 6 month-old *App/Ps1*$^{+/-}$ and *Casp3*$^{-/-}$; *App/Ps1*$^{+/-}$ mice. Mean and S.D. are shown. p-values were calculated with unpaired two-tailed t-tests. n=6 for *App/Ps1*$^{+/-}$ mice and n=4 for *Casp3*$^{-/-}$; *App/Ps1*$^{+/-}$ mice.

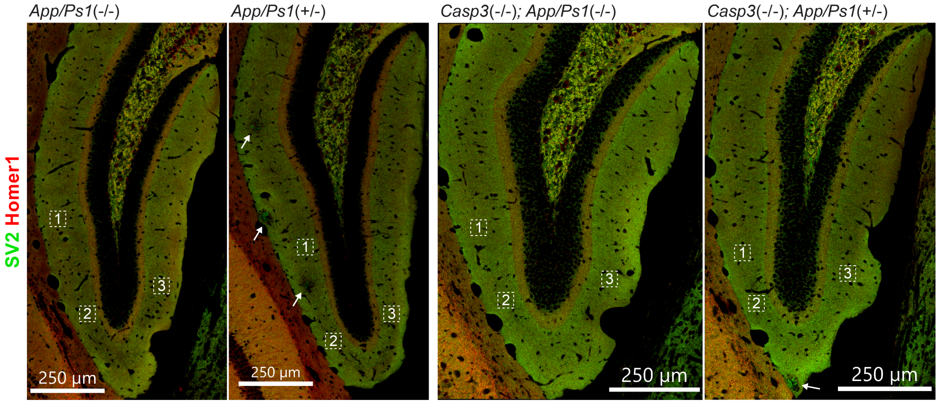


**Figure S15. Selection of field of interest for synapse loss analysis.** To quantify synapse density, three regions of interest from each dentate gyrus on either side of the brain were chosen for each animal (total of 6 regions per animal). Regions of interest in one dentate gyrus of one animal from each genotype group are highlighted with dotted squares in the overviews above. High resolution images of SV2 (in green) and Homer1 (in red) stains were acquired for each region of interest and used for analysis. In sections from animals overexpressing mutant APP and PS1, we avoided choosing regions of interest from areas surrounding amyloid plaques (highlighted with arrows).

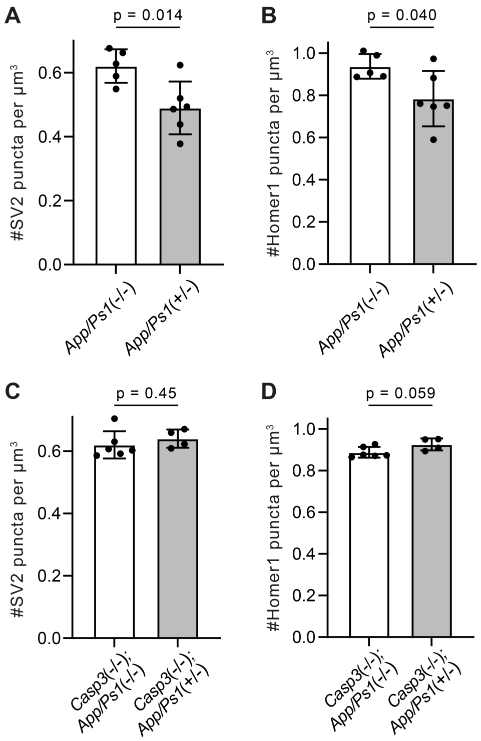


**Figure S16. Additional analysis of synapse density in APP/PS1 mouse lines.** (**A**-**D**) Quantification of presynaptic (**A** and **C**) and postsynaptic (**B** and **D**) puncta densities in the dentate gyrus of APP/PS1 mice in the caspase-3 wildtype (**A** and **B**) or deficient (**C** and **D**) background. Mean and S.D. are shown. p-values were calculated from unpaired two-tailed t-tests. n=5 for *App/Ps1*$^{-/-}$ mice, n=6 for *App/Ps1*$^{+/-}$ mice, n=6 for *Casp3*$^{-/-}$; *App/Ps1*$^{-/-}$ mice, and n=4 for *Casp3*$^{-/-}$; *App/Ps1*$^{+/-}$ mice.

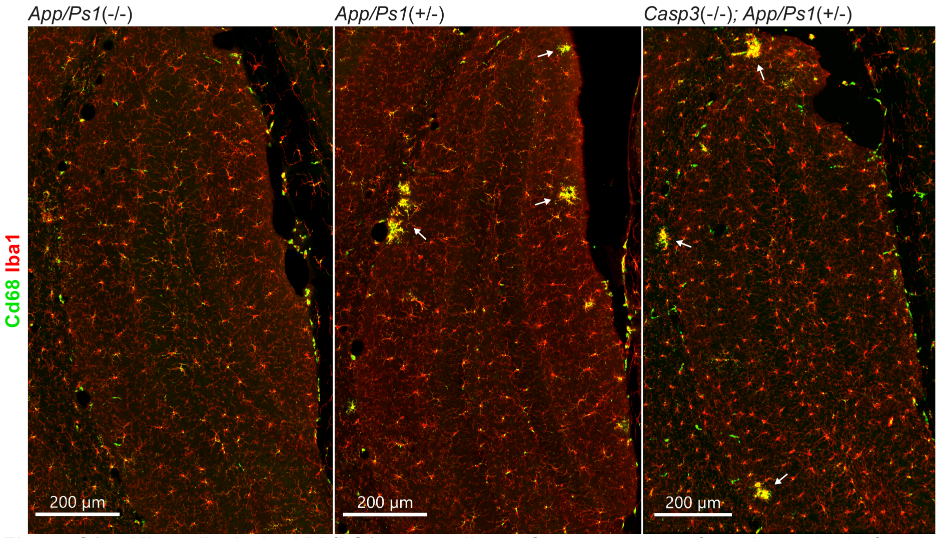


**Figure S17. Microgliosis in APP/PS1 mouse lines.** Coronal sections from 6 month-old female *App/Ps1*$^{-/-}$ (left), *App/Ps1*$^{+/-}$ (middle), and *Casp3*$^{-/-}$; *App/Ps1*$^{+/-}$ (right) mice were stained for Iba1 (red, to detect microglia) and Cd68 (green, as a microglia activation marker). Dentate gyrus regions are shown in the overviews above. Clusters of reactive microglia surrounding amyloid plaques are highlighted with arrows.

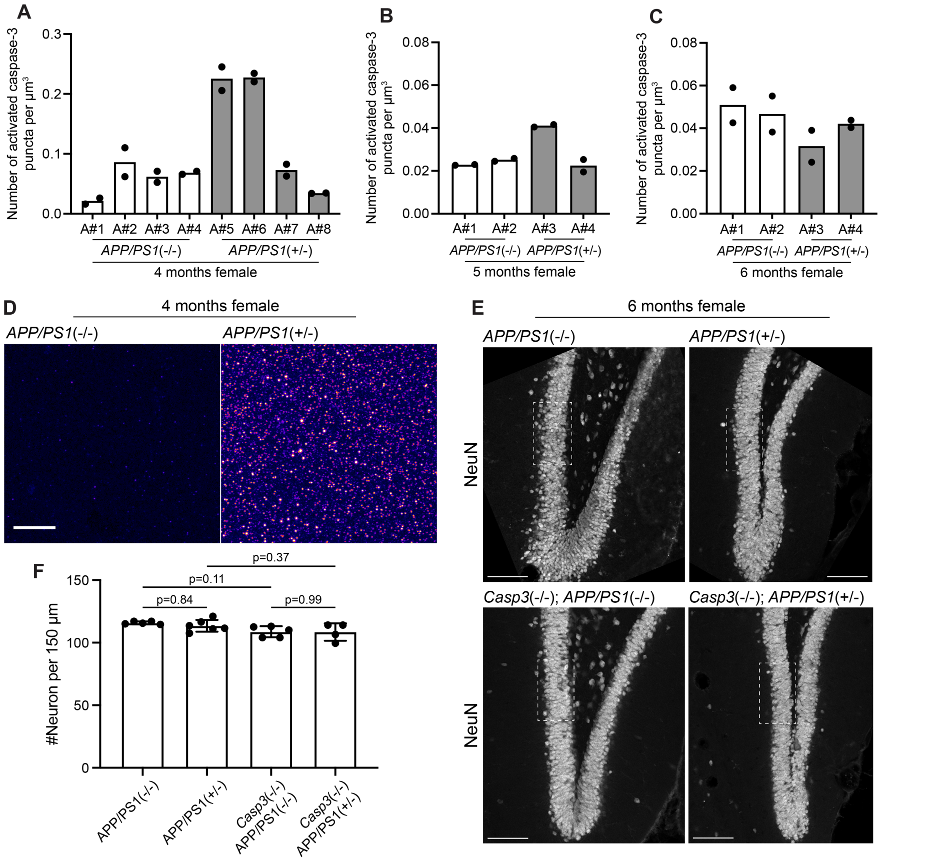


**Figure S18. Aβ-induced caspase-3 activity in APP/PS1 mice.** (**A**-**C**) Density of activated caspase-3 puncti in the dentate gyrus of 4 month-old (**A**), 5 month-old (**B**), and 6 month-old (**C**) female *App/Ps1*$^{-/-}$ and *App/Ps1*$^{+/-}$ mice. Each column represents one animal, and each dot represent quantification from one dentate gyrus (either from left side or right side). Two out of four 4-month-old *App/Ps1*$^{+/-}$ mice showed robustly upregulation of caspase-3 activity in the dentate gyrus (**A**). No upregulation of caspase-3 activity was observed in 5 month-old or 6 month-old mice. (**D**) Images showing elevated caspase-3 activity in the molecular layer of the dentate gyrus of a 4 month-old female *App/Ps1*$^{+/-}$ mouse (A#5 in **A**). Images were set to the same intensity contrast. Upregulated caspase-3 activity remained in a punctate pattern and no apoptotic cell was observed. Scale-bar represents 10 μm. (**E**) Representative images of the dentate gyrus in 6 month-old female *App/Ps1*$^{-/-}$, *App/Ps1*$^{+/-}$, *Casp3*$^{-/-}$; *App/Ps1*$^{-/-}$, and *Casp3*$^{-/-}$; *App/Ps1*$^{+/-}$ mice that were stained with an anti-NeuN antibody to visualize neurons. The fields of interest used to quantify neuron density are highlighted with dotted rectangles. Scale-bars represent 100 μm. (**F**) Neuron density in the granule cell layer of the dentate gyrus in the four groups of mice shown in **E**. Neither caspase-3 deficiency nor APP/PS1 overexpression caused significant changes in neuron density. Mean and S.D. are shown. p-values were calculated from Tukey's multiple comparisons test. n=5 for *App/Ps1*$^{-/-}$ mice, n=6 for *App/Ps1*$^{+/-}$ mice, n=5 for *Casp3*$^{-/-}$; *App/Ps1*$^{-/-}$ mice, and n=4 for *Casp3*$^{-/-}$; *App/Ps1*$^{+/-}$ mice.
