## Supplementary material for "Activity-dependent synapse elimination requires caspase-3 activation": methods

**Material and methods**

**Mouse lines**

The *Casp3*^-/-^ mouse line was a generous gift from Dr. Richard Flavell. This mouse line was received as cryopreserved sperm and was rederived using C57Bl/6J oocytes. The resulting colony was maintained by continuous backcrossing to C57Bl/7J mice. We verified that mice in our *Casp3*^-/-^ colony were congenic to the C57Bl/6J background (greater than 99% identity by SNP analysis). It is important to note that regular backcrossing is necessary to maintain phenotype stability in *Casp3*^-/-^ mice. A small fraction of *Casp3*^-/-^ mice displayed varying degrees of hydrocephalus. These animals were excluded from our analyses. *Cx3cr1-Gfp* mice (stock number 005582) were obtained from the Jackson laboratory. *Aldh1l1-Gfp* transgenic mice (stock number 011015-UCD) were obtained from Mutant Mouse Resource and Research Centers (MMRRC) as cryopreserved sperm and rederived using C57Bl/6J oocytes. *App/Ps1* mice were obtained from the Jackson laboratory (stock number 034832). Mice carrying *App/Ps1* transgenes were given DietGel^®^ 76A (ClearH_2_O, 72-07-5022) and were euthanized at the age of 6 months if not used in experiments. For experiments associated with Figure 1 and Figure 2, C57Bl/6NCrl mice were obtained from Charles Rive Laboratories (strain code 027). For experiments associated with Figure 6, C57Bl/6J mice were obtained from the Jackson Laboratory (stock number 000664). All animal procedures were carried out according to IACUC approved protocols.

**Reagents**

Reagents were obtained from the following sources: isoflurane liquid (NDC 50989-150-15) was from Vedco; phosphate buffered saline without calcium and magnesium (PBS^-^, BP2438) was from Fisher Scientific; bicuculline (0130) and cyclothiazide (0713) were from Tocris Bioscience; PBS with calcium and magnesium (PBS^+^, SH30264) was from Cytiva; dimethyl sulfoxide (DMSO, D2650), bovine serum albumin (BSA, A9647), triton X-100 (T8787), Thioflavin S (T1892) were from Millipore-Sigma; paraformaldehyde powder (19200) was from Electron Microscopy Science; RPMI 1640 medium (11875093), DMEM medium (11965092), fetal bovine serum (FBS, 16000044), penicillin/streptomycin (pen/strep, 15140122), CD Hybridoma medium (11279023), GlutaMAX supplement (35050061), protein A/G agarose beads (20421), protein A/G IgG binding buffer (54200), IgG elution buffer (21004), 1M Tris-HCl pH 8.0 (15568025), 10% normal goat serum (50062Z), ProLong^TM^ Gold mounting medium (P36934), Alexa Fluor 546 succinimidyl ester (A20002), cholera toxin subunit beta(CTB) conjugated to AlexaFlour (AF) 488, AF555, AF594 and AF647 (C22841, C22842, C22843, C34778), goat anti-rabbit secondary antibodies conjugated to AF488plus (A32731), AF555plus (A32732), AF594plus (A32740), and AF647plus (A32733), goat anti-chicken secondary antibody conjugated to AF488plus (A32931), goat anti-guinea pig secondary antibody conjugated to AF488 (A11073) and AF594 (A11076) were from ThermoFisher Scientific; anti-cleaved caspase-3 antibody (9661) was from Cell Signaling Technologies; anti-GFP antibody (GFP-1010) was from Aves Lab; anti-MAP2 antibody (NB300-213) was from Novus Biologicals; anti-NeuN antibody (266004) was from Synaptic Systems; anti-RBPMS antibody (1832-RBPMS) was from PhosphoSolutions; anti-Iba1 (ab178846) and anti-Homer1 (Ab97593) antibodies were from Abcam; anti-Cd68 antibody (MCA1957) was from Bio-Rad; hybridoma line producing anti-SV2 antibody (clone name SV2) was from DSHB.

**DNA cloning and AAV preparation**

To make the pAAV-hSyn-mTurquoise2 plasmid, we ordered the DNA fragment for mTurqouise2 (*1*) as a Gblock from IDT and cloned it via isothermal assembly into a linearized AAV expression plasmid (derived from pAAV-Syn-iCre; Addgene #122518) under the control of human synapsin promoter. To make the pAAV-hSyn-TeTxLC-P2A-mTurqouise2 plasmid, the tetanus toxin light chain (TeTxLC) sequence along with the P2A sequence (2A peptide from porcine teschovirus-1 polyprotein) at the 3’ end was obtained via PCR from Addgene plasmid #129102 and fused in-frame with the mTurquoise2 DNA sequence in the same plasmid backbone as the pAAV-hSyn-mTurquoise2 plasmid. Note that in the main text, we used pAAV-hSyn-TeTxLC to refer to pAAV-hSyn-TeTxLC-P2A-mTurquoise2 because AAV carrying this plasmid induced very low levels of mTurquoise2 expression *in vivo*. When we needed endogenous florescence labeling, we chose to premix AAV carrying pAAV-hSyn-TeTxLC-P2A-mTurquoise2 with other AAVs expressing fluorescent proteins (at 1:1 ratio).

The pAAV-hSyn-tdTomato plasmid was cloned and provided by Janelia Viral Tools. The pAAV-CAG-GFP plasmid was previously made by Svoboda lab and is available as Addgene #28014.

The recombinant AAVs were prepared by Viral Tools of Janelia Core Services. Briefly, two days before transfection, 293T cells were seeded in three 150 mm T/C dishes at 1 x 10^7^ cells/dish in DMEM medium supplemented with 10% FBS and cultured at 37 °C with 5% CO_2_. The cells were transfected with 84 µg of pAAV plasmid, DJ capsid plasmid (*2*), and pHelper plasmid (Agilent 240071) at a ratio of 5:2:3 in polyethylenimine. After 6 to 8 hours, the cells were replenished with serum-free DMEM and further incubated for 3 days. The recombinant AAVs were collected from both the cells and supernatant and purified by two rounds of continuous cesium chloride density gradient centrifugation, dialyzed, concentrated, and filter sterilized. Titer (genome copies per ml) was determined by digital PCR with primers targeting the ITR. For all in vivo experiments a titer of at least 10^13^ genome copies/ml was used.

**Intraocular injections**

*In utero* intraocular injections were performed as previously described (*3*) with modifications. Briefly, pregnant mouse females with embryos at E15 gestation were anesthetized by 3% isoflurane inhalation, abdomens shaved, eye lube applied to each eye, then transferred to a nose cone and set to maintenance level of 2% isofluorane inhalation anesthesia. Warm fluids (lactated Ringer's solution + 5% dextrose, Strategic Applications Inc.), Buprenorphine (0.1 mg/kg, Butler Animal Health), and Ketoprofen (5 mg/kg, Covetrus) were administered subcutaneously just prior to starting surgery. The animal was placed in dorsal recumbency, and the abdomen was cleansed using alternating swabs of 70% ethanol and 4% Chlorhexidine Gluconate (Molnlycke Health Care).  A sterile drape was placed over the abdomen and using aseptic technique, the abdominal skin and wall were cut at midline, before gently removing the uterus from the abdomen. While keeping the uterus/embryos hydrated with warm saline sterile solution, the embryos were gently positioned with ring forceps to hold each embryo securely for injection.  A previously filled glass pipette (Drummond Scientific), beveled to 30-40 μm at the opening was utilized to deliver constructs utilizing a pre-programmed Nanoject III microinjector (Drummond Scientific).  200-250 nl of AAVs prep was injected through the uterine wall and into each eye at a speed of 50nl per second. To confirm proper targeting within the eye, the AAV solution was pre-mixed with 0.05% Fast Green (Sigma Aldrich). After all injections were completed, the uterus/embryos were carefully placed back into the abdomen and the abdomen was flushed with warm sterile saline. The abdominal wall and skin were sutured, and Marcaine (Hospira) was applied subcutaneously below the skin incision.  A small drop of Vetbond (3M) was applied to seal the skin incision. The animal was then removed from the anesthesia apparatus and allowed to recover on a heating pad before being placed back into a clean cage.

For intraocular injection at P4 and P9, pups with desired genotypes were anesthetized in an induction chamber filled with 2.5% - 3% isoflurane using a calibrated vaporizer. After the onset of anesthesia, isoflurane is reduced to 1.25%, and the mouse was removed from the chamber and placed under a dissection microscope. Proparacaine ophthalmic solution (0.5%, Sandoz, NDC 61314-016-01) was applied to each eye to be injected. A re-breathing nose cone was used to deliver 1.25% – 2% isoflurane throughout the procedure. A gentle blunt dissection was performed at each eyelid junction utilizing closed No.5 Dumont forceps (Fine Science Tools) to create a 2-3 mm opening. A 33-gauge sharp beveled needle attached to a Hamilton microliter syringe (Hamilton Company) was used to draw up CTB-AF solutions. Just prior to injecting, the eye was proptosed and held securely in place by the Dumont forceps in preparation for injection. While holding the eye securely in place, the sclera was punctured with the Hamilton needle at the level of the ora serrata to inject 2.5 µL (for P9) or 2 µL (for P4) of desired CTB-AF solution into each eye. After the injection, eye lube was applied, the eye was allowed to move back into the socket, and the eyelids were gently pressed close to protect the eye. Pups were allowed to recover from anesthesia on a heated pad and put back into the cage with the dam.

**Monoclonal antibody production and labeling**

SV2 hybridoma cells were thawed and seeded in RPMI 1640 medium containing 10% FBS and 1% Pen/Strep at 0.3 million cells/ml and cultured in an incubator at 37 ^°^C with 5% CO_2_. After reaching 1x10^6^ viable cells/ml density, the cells were diluted 1:2 by adding an equal volume of serum free CD Hybridoma medium supplemented with 4% GlutaMax^Tm^ and returned to the incubator. This dilution process was repeated 2 more times so that the cells reached 1x10^6^ viable cells/ml density in a mixed medium containing ~1% FBS. Cell viability obtained was at least 90%. The cells were then spun down at 150xG for 10 minutes and resuspended in complete CD Hybridoma medium at 5x10^5^ cells/ml density. The cells were further expanded by adding more complete CD Hybridoma medium when cells reach 1x10^6^ cells/ml density. Once the desired total volume is reached (typically 100 ml), cells were cultured for 2 weeks without medium change.

Culture supernatant containing SV2 antibody was cleared of hybridoma cells by centrifuging at 1000xG for 10 min. Cleared supernatant was diluted 1:1 with IgG binding buffer and passed through 1 ml of protein A/G agarose bead slurry (ThermoFisher 20421, corresponding to 500 ul of settled resin) packed in a gravity flow column. The column was washed with IgG binding buffer, and SV2 antibody was eluted in five 1 ml fractions using IgG elution buffer and immediately neutralized with 10% volume of 1M Tris-HCl, pH 8.0. Fractions with highest antibody concentration were pooled, desalted with Zeba columns (ThermoFisher 89891), and concentrated to ~1 mg/ml using Amicon centrifuge filters (Millipore UFC801024). After adding glycerol (25% final), antibodies were flash frozen in liquid nitrogen and kept at -80 ^°^C.

To label the primary SV2 antibodies, we used Alexa Fluor 546 succinimidyl ester. Briefly, we buffer-exchanged and concentrated SV2 IgG to 3 mg/ml using Amicon filters (UFC501024 Millipore Sigma) in phosphate buffered saline that was brought to pH 8.3 with 100 mM NaHCO_3_. Dimethyl sulfoxide-dissolved Alexa Flour 546 succinimidyl ester was gently mixed with the SV2 IgG at 10:1 molar ratio and incubated for two hours at room temperature on a rotating shaker. Alexa Fluor 546 labeled IgG were purified from unreacted succinimidyl esters with the aid of PD MiniTrap Sephadex G-25 column (Cytiva 28918007) in PBS^-^ by gravity flow following the manufacturer’s instructions. The degree of labeling was determined by spectrophotometry on Nanodrop One (ThermoFisher) by accounting for the Alexa Fluor 546 extinction coefficient at 554 nm and the corrected protein absorbance at 280 nm. Typically, a degree of labeling of 8-9 mols of dye per mol of protein was obtained. Labeled SV2 were aliquoted and flash-frozen in 25% glycerol for long term storage at -80 ^°^C at a final concentration of approximately 0.75 mg/ml.

**Immunohistochemistry**

The mice were deeply anesthetized in a chamber filled with saturating isoflurane gas and transcardially perfused first with PBS^-^ and then with paraformaldehyde (PFA) solution (4% w/v in 20.3 mM NaH_2_PO_4_ and 79.7 mM Na_2_HPO_4_, pH = 7.4). The brains were removed and post-fixed in 4% PFA solution overnight at 4 ^°^C with gentle shaking. Fixed brains were washed 3x15 minutes in PBS^-^ at room temperature and stored in PBS^-^ at 4 ^°^C. For long term storage, fixed brains were cryoprotected in 30% Sucrose (w/v in PBS^-^) for 72 hours and frozen in Tissue-Tek^®^ O.C.T. compound (Sakura) and stored at -80 ^°^C.

Brains stored in PBS^-^ were imbedded in 4% agarose, and coronal sections (50 um thick for mice younger than P10 or 40 μm thick for adult mice) were prepared using a vibratome (Leica VT 1200S). For brains frozen in O.C.T. compound, coronal sections were prepared using a cryostat (Leica 3050S). Sections were floated in 24-well plates in PBS^-^ and stored at 4 ^°^C. For antibody staining, free-floating sections were first permeabilized/blocked in 10% normal goat serum containing 0.3% (w/v) triton X-100 for 1 hour. The sections were then incubated with primary antibodies diluted in staining buffer (PBS^+^ containing 2% BSA w/v and 0.3% triton X-100) at 4 ^°^C overnight with gentle agitation. Dilutions of primary antibodies were: anti-cleaved caspase-3, 1:500; anti-GFP, 1:2000; anti-MAP2, 1:1000; anti-NeuN, 1:500; anti-Iba1, 1:500; anti-Cd68, 1:500; anti-SV2, dilute to 2.5 μg/ml; anti-Homer1, 1:400. The sections were then washed 3x15 minutes in washing buffer (PBS^+^ containing 0.1% triton X-100) and stained with appropriate fluorophore-conjugated secondary antibodies diluted in staining buffer for 3 hours at room temperature with gentle agitation. Secondary antibodies were always used at 1:500 dilution. The sections were washed 3x15 minutes in washing buffer, rinsed 3x5 minutes in PBS^+^, and mounted onto glass slides (Fisher, 12-550-15). After air-drying completely, mounted sections were immersed in ProLong^TM^ Gold and coverslipped (Fisher, 12-541-024). The sections were imaged after 24 hours of curing at room temperature. The sections were protected from light during the entire process.

To prepare whole-mount retinae, eyes were removed from PFA perfused mice and placed in PBS^-^. The cornea was punctured with a #11 scalpel, and several anterior-to-posterior cuts were made on the sclera starting from the puncture. Intact retinae were isolated by tearing and peeling off the sclera along the cuts. 4 to 5 radial cuts were made on each retina to flatten the tissue. Isolated retinae were post-fixed for 15 minutes in 4% PFA solution at room temperature with gentle agitation and washed for 3x5 minutes in PBS^-^ at room temperature with gentle agitation. Free-floating retinae were then stained with primary (anti-RBPMS antibody, used at 1:200 dilution) and secondary (goat anti-guinea pig Alexa Fluor 594plus) antibodies and mounted onto glass slides as described above.

For staining of amyloid-β (Aβ) plaques, a 1% Thioflavin S stock solution (w/v, dissolved in ddH_2_O) was diluted 100-fold in 50% ethanol (prepared by mixing pure ethanol and ddH_2_O at 1:1 ratio) to create a 0.01% (w/v) working solution. The stock and working solutions were discarded after each use. Free-floating brain sections were dehydrated by washing 2x5 minutes in 50% ethanol, stained in 0.01% Thioflavin S solution for 8 minutes, washed for 2x5 minutes in 50% ethanol, then re-hydrated by washing 3x5 minutes in PBS^+^. All staining steps were performed at room temperature with gentle agitation. The sections were protected from light throughout the procedure. Stained sections were then mounted and cured as described above.

**Measuring synapse inactivation-induced caspase-3 activity**

For experiments associated with Figure 1 and Figure S4, E15 C57Bl/6NCrl mice were injected in the right eyes either with AAV carrying hSyn-mTurquoise2 (control group), or with a 1:1 mixture of AAV carrying hSyn-mTurquiose2 and AAV carrying hSyn-TeTxLC-P2A-mTurquoise2 (synapse inactivation group). At the age of P5, brains of both groups of mice were harvested and sectioned as described above. To quantify caspase-3 activity in the dLGN and the localization of activated caspase-3 relative to TeTxLC-expressing RGC axons (Figure 1), sections were stained with anti-cleaved caspase-3 antibody (using goat anti-rabbit AF594plus as the secondary) and anti-GFP antibody (using goat anti-chicken AF488plus as the secondary). Alternatively, sections were co-stained with anti-cleaved caspase-3 (using goat anti-rabbit AF647plus as the secondary) and anti-MAP2 antibody (using goat anti-rabbit AF488plus as the secondary) (Figure S4A), or with anti-cleaved caspase-3 (using goat anti-rabbit AF594plus as the secondary) and anti-NeuN antibody (using goat anti-rabbit AF488plus as the secondary) (Figure S4C). Immunostaining was performed as described above.

For experiments associated with Figure 2, E15 C57Bl/6NCrl mice were injected in the left eyes with AAV carrying hSyn-tdTomato and in the right eyes with AAV carrying CAG-eGFP (control group), or in the right eyes with AAV carrying hSyn-mTurquoise2 and AAV carrying hSyn-TeTxLC-P2A-mTurquoise2 (single inactivation group), or in the left eyes with AAV carrying hSyn-tdTomato and AAV carrying hSyn-TeTxLC-P2A-mTurquoise2 and in the right eyes with AAV carrying CAG-eGFP and AAV carrying hSyn-TeTxLC-P2A-mTurquoise2 (dual inactivation group). At the age of P5, brains of all three groups of animals were harvested, sectioned, and stained with anti-cleaved caspase-3 antibody and goat antirabbit AF647plus secondary antibodies as described above.

To quantify caspase-3 activity in the dLGN (Figure 1B-E), confocal images of dLGNs were acquired on a Leica Stellaris 8 microscope using a 10x/0.4 NA dry objective (Leica HC PL Apo CS2 10x/0.4 dry). The channels were acquired sequentially (line-by-line) with the anti-caspase-3 first and anti-mTurquoise2 second. A 590 nm laser and emission read of 596 nm – 748 nm was used for the anti-cleaved caspase-3 channel, and 440 nm and 488 nm lasers and an emission read of 457 nm – 579 nm were used for anti-mTurquoise2 channel. The imaging parameters (laser power, pixel dwell time, signal averaging or accumulation) were set to maximize signal-to-background values while avoiding detector (SiPM type Power HyD S) saturation. Photon detection was set in counting mode and acquired as 12-bit images at 227 nm x 227 nm pixel size. All datasets used for measurement of caspase-3 activity were acquired as Z-stacks (7 slices spaced 2.4 μm apart). To measure the total amount of active caspase-3 signal in the 3D stacks of dLGN we used the Labkit (*4*) Fiji plugin to generate and train a pixel classifier through Imaris v. 10 (Oxford Instruments). The same pixel classifier and surface creation settings were applied to data from both the control and the synapse inactivation groups to generate surface objects of active caspase-3 signal. All active caspase-3 objects found within the dLGN area (manually delineated based on mTurquoise2 signal) were selected, and the sum of voxel intensities the active caspase-3 objects was calculated and divided by the area of dLGN. The final values plotted for each mouse are averages of two consecutive sections.

To image active caspase-3 signals and TeTxLC-expressing RGC axons at high resolution (Figure 1G), we used a Leica Stellaris 8 confocal microscope and a 63x/1.4 NA oil objective (Leica HC PL APO 63x/1,40 OIL CS2) in “Lightning” mode (in LasX 4.6) set at high resolution grade, which reduces the pinhole size to 0.5 Airy units. Laser and emission settings were identical to those used the previous section that described quantification of active caspase-3 signal in mTurquoise2-labeled dLGNs. 3D image stacks (31.5 um x 31.5 um x 12 um) were acquired at high voxel density (43 nm x 43 nm x 204 nm) in photon counting mode and 4x signal accumulation. Subsequently, the images were processed by iterative deconvolution using a theoretical point spread function informed by the imaging parameters of the microscope and constrained by an “Adaptive” strategy that accounts for local background and signal-to-noise ratio information.

Confocal images of MAP2 and active caspase-3 signals in dLGN (Figure S4A) were similarly imaged on a Leica Stellaris 8 microscope with a 63x/1.4 NA oil objective (Leica HC PL APO 63x/1,40 OIL CS2). A 653 nm laser line and an emission read of 662 nm – 829 nm were used for caspase-3 channel, and a 499 nm laser line and an emission read of 504nm – 549 nm were used for Map2 channel. The channels were acquired simultaneously at voxel densities of 65 nm x 65 nm x 299 nm. NeuN and active caspase-3 signals in dLGN (Figure S4C) were imaged using a 20x/0.7 NA water objective (Leica HC PL APO 20x/0,7 IMM CORR CS2). A 590 nm laser and an emission read of 595 nm – 750 nm were used for imaging caspase-3 and a 499 nm laser with a 509 nm - 595 nm emission read was used for the NeuN signal. The channels were acquired sequentially (active caspase-3 first) at pixel densities of 142 nm x 142 nm.

Measuring overlap of eye-specific territories in dLGN

For comparison between Casp3^+/+^ and Casp3^-/-^ animals (Figure 3), Casp3^+/-^ males were bred to Casp3^+/-^ females to generate Casp3^+/+^ and Casp3^-/-^ littermate pups. Pups were genotyped at birth. Before intraocular injection, lyophilized CTB-AF488 and CTB-AF594 powders were reconstituted in 1% DMSO in PBS^-^ to a final concentration of 5 mg/ml. CTB solutions were aliquoted and stored at -20 °C for later use. Thawed CTB solutions were discarded after each surgery. Intraocular injection of CTB-AF solutions was performed at the age of P9 as described above. CTB-AF488 was injected into left eyes and CTB-AF594 into right eyes. 24 hours after the injection, brains of P10 pups were harvested, and 50 um thick coronal sections prepared as described above.

Cured sections were imaged on a Zeiss AxioObserver Z.1 microscope equipped with an LSM 880 Airy scan detector and an EC Plan-Neofluar 10x objective (NA = 0.3). A 488 nm laser and a 499-544 nm detection window were used for CTB-AlexaFlour488, and a 594 nm laser and a 597-659 nm detection window were used for CTB-AlexaFlour594. Signals from the two channels were acquired sequentially into 16-bit images with a 0.4 µm x 0.4 µm pixel size. Laser power and detector gain settings were maintained at the same level for most images, with occasional adjustment to avoid saturation. One image per dLGN per section was acquired.

To quantify the overlap between contralateral-specific and ipsilateral-specific territories in the dLGN, we selected 7-8 sections per animal that span the majority of left dLGN of and analyzed their images in ImageJ. For each image, a mask for the entire dLGN region and a mask for a background region in the thalamus with no labeling were manually created. Background signal from each channel was estimated as the average signal in the background region and was subtracted from signals in the dGLN region. Then the background-subtracted signals in the dLGN region were normalized to between 0 and 1 for each image and each channel. We picked 7 increasing cutoff thresholds (0.1, 0.125, 0.15, 0.175, 0.2, 0.225, and 0.25), and for each threshold and each channel, signals that are greater than or equal to the threshold were considered real signals. For each animal (corresponding to 7-8 images of left dLGN) and each threshold, we calculated percentage overlap as the ratio between the total number of pixels with real signals in both channels in all images and the total number of pixels in the entire dLGN region of all images. We plotted the percentage overlap as a measurement of segregation of eye-specific territories in dLGN. Similar analyses were performed on right dGLN images yielding similar results.

For overlap analysis presented in Figure S3, E15 mouse embryos were injected in their right eyes either with AAV carrying hSyn-mTurqouise2 (control group) or a 1:1 mixture of AAV carrying hSyn-TeTxLC-P2A-mTurqouise2 and AAV carrying hSyn-mTurquoise2 (inactivation group). The left eyes of both groups of mice were injected with AAV carrying the hSyn-tdTomato. At P8 (when segragation of eye territories is largely complete and thus sufficient for us to validate the effect of TeTxLC on synapse inactivation), brains from both groups were harvested, and 50 μm thick coronal sections were prepared as described above. Only animals that had well labeled dLGNs on both sides were selected for analysis. Mounted and cured sections containing dLGN were imaged in confocal mode on a Leica Stellaris 8 microscope using a 10x/0.4 NA dry objective (Leica HC PL Apo CS2 10x/0.4 dry). A 441 nm laser and a 445-548 nm detection window were used for the mTurquoise2 channel, and a 554 nm laser and a 564-721 nm detection window were used for tdTomato channel. The images were acquired at 8-bit depth with a 0.227 μm x 0.227 μm pixel size (at 1 Airy unit for λ = 580 nm) and at a laser power, pixel dwell time, and signal accumulation sufficient to avoid detector (Power HyD S in counting mode) saturation. One image per dLGN per section was acquired. For each animal, overlap was measured as described above on the left dLGN (the side where mTurquoise2 signal occupied the majority of the dLGN) by averaging from 3 to 5 consecutive sections. The cutoff thresholds used were: 0.050, 0.075, 0.1, 0.125, 0.15, 0.175, 0.2.

For overlap analysis presented in Figure S5, embryos in the control group were injected at E15 with AAV carrying tdTomato construct in the left eyes, and with AAV carrying the CAG-GFP construct in the right eyes. Embryos in the dual inactivation group were injected at E15 with a mixture (1:1) of AAV carrying hSyn-tdTomato construct and AAV carrying hSyn-TeTxLC-P2A-mTurquoise2 construct in their left eyes, and with a mixture (1:1) of AAV carrying the CAG-GFP construct and AAV carrying the hSyn-TeTxLC-P2A-mTurquoise2 construct in their right eyes. At P10, the brains were harvested and sectioned as described above. Confocal images of dLGN were acquired on a Leica Stellaris 8 microscope using a 10x/0.4 NA dry objective (Leica HC PL Apo CS2 10x/0.4 dry). A 488 nm laser line with a 488 nm notch filter and a 485-559 nm detection window were used for the GFP channel, and a 554 nm laser and a 559-701 nm detection window were used for the tdTomato channel in sequential line mode to avoid any cross-excitation (with tdTomato channel acquired first). The images were acquired at 12-bit depth with a 0.248 μm x 0.248 μm pixel size (at 1 Airy unit for λ = 580 nm) and at a laser power, pixel dwell time, and signal accumulation that avoided detector (Power HyD S in counting mode) saturation. One image per dLGN per section was acquired. For each animal overlap was measured as described above on the right dLGN (the side where tdTomato signal occupied the majority of the dLGN) by averaging from 5 consecutive sections.

**Quantification of RGC densities in the retina**

To estimate the density of RGCs in the retina (Figure S6), retinae of P10 *Casp3*^+/+^ and *Casp3*^-/-^ mice were harvested and stained with anti-RBPMS antibody as described above. We imaged 4 large square regions of interest (1.16 mm x 1.16 mm) in each retina on four sides of the optic disc. The images were acquired with a Leica TCS SP8 Laser Scanning Microscope using the 10x/0.4 NA dry objective (Leica HC PL Apo CS2 10x/0.4 dry), a 594 nm laser line, and an emission window of 607 nm to 694 nm, at a pixel sampling of 302 nm x 302 nm. To account for tissue unevenness, we collected a stack of 3 Z-slices (2.4 μm apart). To count RGCs, we sampled 3 subregions of a size of 100 μm^2^ from each large region of interest. The subregions were selected to be at similar distances from the optic disc and in areas devoid of tissue tears. Cells were modeled to spots by fitting them to ellipsoid objects (XY size = 8 μm; Z size = 16 μm, with background subtraction) in Imaris 9.6 (Oxford Instruments). Densities of spots for each region were averaged and reported as number of RGC per 100 μm^2^ of retina.

**Quantification of dLGN neuron densities**

To determine the density of relay neurons in dLGN (Figure S7), we collected and stained 50 μm brain sections with anti-NeuN antibody as described above. Confocal image stacks (Leica Stellaris 8) were acquired with a 10x/0.4 NA dry objective (Leica HC PL Apo CS2 10x/0.4 dry) at a voxel sampling of 210 nm x 210 nm x 2410 nm. For DAPI (i.e. cell nuclei) signal, the 405 nm laser with 430 nm – 550 nm emission was used, while for the NeuN signal, revealed by Alexa594 conjugated IgG, the 590 nm laser with 597 nm – 750 nm emission was used. For each animal, two consecutive coronal histological sections were imaged as Z stacks containing 8 Z-slices that were centered on dLGN. Volumes of NeuN signal were segmented in Imaris 10.2 (Oxford Instruments). To improve the identification of neuronal cell bodies we generated synthetic images from the geometric means of DAPI and segmented NeuN images and applied a spot object finder algorithm (6 μm diameter) to the entire stack. Finally, we selected the spots corresponding to dLGN only and reported the number of spots over the area of dLGN (manually traced for each histological section). Each data point in Figure S7 is an average of dLGN densities from two brain sections of the same animal.

**Whole-cell patch-clamp recording in acute brain slices**

To characterize electrophysiological properties of retinogeniculate synapses in *Casp3*^+/+^ and *Casp3*^-/-^ mice (Figure 4), the mice (aged p28 – p32) were decapitated under deep isoflurane anesthesia, and the brain was removed and quickly transferred to an ice-cold dissection solution containing (in mM): 194 sucrose, 30 NaCl, 2.5 KCl, 1.2 NaH2PO4, 26 NaHCO3, 10 D-Glucose, 1 MgCl2 (pH 7.4, oxygenated with 95% CO2 and 5% O2). Parasagittal brain slices containing the optic tract and dLGN were prepared using a vibratome (Leica VT 1200S) as previously described (*5, 6*). The slices were recovered at 35.5°C in an incubation chamber (BSC-PC, Warner Instrument, USA) filled with artificial cerebrospinal fluid (ACSF) containing (in mM): 124 NaCl, 2.5 KCl, 1.2 NaH2PO4, 26 NaHCO3, 10 D-Glucose, 2 CaCl2, 1 MgCl2 (pH 7.4, oxygenated with 95% CO2 and 5% O2, osmolarity ~310). After ~30 minutes of recovery, the chamber was maintained at room temperature.

All recordings were performed with a MultiClamp 700B amplifier (Molecular Devices) and signals were filtered at 2 kHz and digitized at 20 kHz with via USB-6343 (National Instruments) under the control of WaveSurfer software (https://wavesurfer.janelia.org). All recordings were carried on slices submerged in the recording chamber of an upright microscope (BX61WI; Olympus, Tokyo, Japan) equipped with IR-DIC (infrared-differential interference contrast) microscopy and a water-immersion objective lens (60X, 1.00 NA; Olympus). Slices were maintained under continuous perfusion of oxygenated ACSF at 29-31°C. Patch pipettes were pulled from thin-wall single-barrel borosilicate glass (TW150-6, WPI), resulting in electric resistance of 3 ~ 5 MΩ when filled with an intracellular solution contained the following (in mM): 120 cesium methane sulfonate, 5 NaCl, 10 tetraethylammonium chloride, 10 HEPES, 4 lidocaine N-ethyl bromide, 1.1 EGTA, 4 magnesium ATP, and 0.3 sodium GTP, with pH adjusted to 7.2 with CsOH and osmolality set to ∼290 mOsm.

Whole-cell voltage-clamp recordings of relay neurons in the dLGN were performed as previously described (Figure 4A-C) (*6*). The slices were submerged in ACSF containing 20 uM bicuculline to inhibit GABA_A_ receptors, and a bipolar concentric stimulation electrode (FHC) was placed on the surface of the optic tract to deliver stimuli with intensities ranged from 0 – 100 µA (200us duration) with an inter-trial interval of 40 – 60 s. In the whole-cell configuration (series resistance < 20 MΩ), the membrane potential of the neuron was clamped at -70 mV to measure evoked excitatory postsynaptic current (EPSC) mediated by activation of AMPA receptors, and then at +40 mV to measure evoked EPSC mediated by activation of NMDA receptors. The stimulus intensity was systemically increased until a response to the stimulus was observed, and then was reduced to a previous level where no response was detected. From that level, the stimulus intensity was increased by 0.5 μA steps to recruit single inputs which were putative single fiber (SF) amplitudes. When the EPSC amplitude reached a plateau, it was considered a putative maximum response. If the EPSC amplitude dropped while the stimulation intensity increased, data from that relay neuron was discarded.

To measure miniature excitatory postsynaptic current (mEPSC), the slices submerged in the ACSF containing TTX 1 µM and bicuculine 20 µM to inhibit spontaneous action potentials and GABA_A_ receptors. The membrane potential was clamped at -70 mV and synaptic current was recorded for 180 s. After the recoding, -9 pA threshold was used to detect mEPSC.

To measure paired pulse ratio (PPR) (Figure 4D-E), the slices were submerged in the ACSF containing 20 uM bicuculline and 50 uM cyclothiazide to inhibit GABA_A_ receptors and AMPAR desensitization, and the stimulation electrode was placed on the optic tract. In the whole-cell configuration, the membrane potential was clamped at -70 mV. After confirming the stimulation intensity that evoked the maximum response, two consecutive electrical stimuli of that intensity (300 µs duration) were delivered at 50, 150, 250, 500, and 1000 ms inter-stimulus intervals. Each pair of stimuli constitutes one trial, and two consecutive trials were separated by an interval of 40 s. For each inter-stimulus interval, two trials were performed and averaged. PPR was calculated as the maximum amplitude of the second EPSC over the maximum amplitude of the first EPSC. If in any trial except for the first trial the first EPSC dropped significantly in amplitude compared to the first EPSC of the first trial, data from that dLGN relay neuron was discarded.

**Measuring microglia- and astrocyte-mediated synapse engulfment *in vivo***

To measure microglia-mediated synapse engulfment *in vivo* (Figure 5 and Figure S10), we bred *Casp3*^-/-^ males to *Casp3*^+/-^; *Cx3ct1-Gfp^+/-^* females to generate *Casp3*^-/-^; *Cx3ct1-Gfp^+/-^* and *Casp3*^+/-^; *Cx3ct1-Gfp^+/-^* littermate pups. To measure astrocyte-mediated synapse engulfment *in vivo* (Figure S11), we bred *Casp3*^-/-^ males to *Casp3*^+/-^; *Aldh1l1-Gfp^+/-^* to generate *Casp3*^-/-^; *Aldh1l1-Gfp^+/-^* and *Casp3*^+/-^; *Aldh1l1-Gfp^+/-^* littermate pups. P4 pups with desired genotypes were injected with 2 µL of CTB-AF555 in left eyes and 2 µL of CTB-AF647 in right eyes as described above. 24 hours later, the brains were harvested, sectioned, and mounted as described above.

To analyze engulfment of synapses, the sections were imaged using a Leica TCS SP8 Laser Scanning Microscope equipped with an HC Plan Apo CS2 40x (NA = 1.3) oil objective and a White Light Laser Unit. The 3D multichannel stacks with a voxel size of 0.075 x 0.075 x 0.312 µm for microglia samples or 0.048 x 0.048 x 0.2 µm for astrocytes samples were acquired as 16-bit images with 4x line averages for microglia or 2x for astrocytes at 400 Hz scanner speed. The stacks were set to collect a volume of 20 micrometers in the Z dimension and 387.5 micrometers in XY dimensions for microglia or 288 μm in XY for astrocytes. The pinhole was set at 1 Airy unit for 520 nm wavelength.  The excitation lasers and detection windows were set to minimize crosstalk and bleed-through as well as the saturation of detectors while simultaneously imaging the three channels. A 478 nm laser and a detection window of 488 nm – 537nm were used for the GFP channel, a 550 nm laser and a detection window of 596 nm – 629 nm were used for CTB-AF555, and a 651 nm laser and a detection window of 665 nm to 750 nm were used for CTB-AF647. Imaris 9.6 (Oxford Instruments) was used to segment microglia, astrocytes, ipsilateral RGC axon terminals, and contralateral RGC terminals from the 3D stacks using the Surface Creation module with background subtraction. For segmentation of RGC axon terminals and astrocytes, all parameters used were left at default values (auto). For segmentation of microglia, we set the threshold manually at the highest inflection curve in the histogram of voxel intensities, which in our experience provided a good trade-off for good segmentation of cell bodies and major processes while keeping the adjustment consistent across different image datasets. A background subtraction ball size of 1 µm was used for microglia, a ball size of 1.4 µm was used for astrocytes, and a ball size of 0.5 µm was used for AF555 and AF647 labeled axonal materials. The segmented signal of AF555 and AF647 was each converted to a binary mask of 0 and 1 voxels. Finally, the sum of the segmented volume (i.e. sum of ones) of AF555 and or AF647 in each microglia or astrocyte was either directly plotted or normalized to the microglia or astrocyte volume and plotted. To control the inherent signal variability across LGN, only microglia and astrocytes present in the ipsilateral region of the 3 middle sections of the left dLGN of each mouse were included in the analysis. Also, only microglia or astrocytes whose soma and proximal processes were contained in the volumetric stack were included in the final analysis. All littermate pups of the same genotype were pooled together.

**Measuring activity-dependent engulfment of synapses by microglia**

To investigate the role of caspase-3 in activity-dependent synapse elimination (Figure 6 and Figure S12), we used C57Bl/6J wildtype mice as controls and bred *Casp3*^-/-^ males with *Casp3*^+/-^ females to generate *Casp3*^-/-^ mice. Note that C57Bl/6J and *Casp3*^-/-^ mice used in this experiment are not littermates and should not be directly compared with each other. C57Bl/6J and *Casp3*^-/-^ embryos were injected at E15 in their right eye either with AAV carrying hSyn-TeTxLC-P2A-mTurqouise2 or AAV carrying hSyn-mTurquoise2 as described above. To label the RGC axons, at P4, the pups were injected with 2 µL of CTB-AF555 in left eyes and 2 µL of CTB-AF647 in right eyes as described above. At P5, the brains were harvested and sectioned. Two 50 μm sections that spanned the central part of the dLGN were stained with anti-Iba1 antibody and goat anti-rabbit AF488 secondary (as described above) to label microglial cell bodies.

Mounted and cured sections were then imaged on Leica Stellaris 8 confocal microscope using a 63x/1.4 NA oil objective (Leica HC PL APO 63x/1,40 OIL CS2). Four stitched 3D-stacks were acquired for each left dLGN (the side where CTF-AF647 and mTurquoise2 occupied the majority of the dLGN) to generate a 3D volume of 366 μm x 366 μm in XY and approximately 20 μm in the Z dimension, with a voxel size of 64 nm x 64 nm x 299 nm. The channels were acquired sequentially: first using a 499 nm laser with an emission window of 507-556 nm together with a 653 nm laser with an emission window of 663-750 nm for Iba1-AF488 and CTB-AF647 signal, respectively, then using a 553 nm laser with an emission window of 558-663 nm for the CTB-AF555 signal. Images acquired were collected as 12-bit datasets, and care was taken to set the laser power, signal accumulation, and pixel dwell time to avoid detector saturation. Only animals whose dLGN areas were fully labeled with both AF555 and AF647 as well as the mTurquoise signals were imaged and analyzed. Animals in which CTB signal appeared widespread throughout the brain parenchyma were excluded from the analysis as these labeling patterns were likely due to surgical mistargeting or issues with tissue integrity.

To segment volumes corresponding to microglia and CTB-AF555 and CTB-AF647 labeled RGC axon terminals, we utilized the Surface Creation module with machine learning segmentation of Imaris 10.1.1 (Oxford Instruments). We first trained for several rounds a pixel classifier for each channel using two independent image files and saved the classifiers along with the other settings (smoothing grain size of 100 nm and discarding volumes smaller than 10 voxels) as “Favorite Creation Parameter”. Subsequently, the same Favorite Creation Parameters were applied to all data sets, with the rare instances when microglia segmentation was slightly tuned via the same pixel classier to better capture some of the cell border areas. Similar to the previous analyses (Figure 5), the segmented signals of CTB-AF647 and CTB-AF555 were converted to binary images (0 or 1 voxels), and the total volume of CTB-AF647- or CTB-AF555-positive voxels in each microglia was extracted, converted to fraction of the microglia volume, and plotted. Only the cells whose body and major processes were bound within the 3D stack and present within the middle area of dLGN (where the ipsilateral signal is present) were included in the analysis. Data from the same genotype and treatment were pooled together. Either one or two sections were analyzed for each animal.

**Measuring microglia activation**

To measure if lack of caspase 3 causes microglia activation (Figure S9), we bred *Casp3*^+/-^ males with *Casp3*^+/-^ females to generate *Casp3*^-/-^ and Casp3^+/+^ littermates. After genotyping, at P5, the brains were harvested and sectioned coronally as described above. Two 50 μm sections that spanned the central part of the dLGN were stained with anti-Iba1 antibody (detected by goat anti-rabbit AF488 secondary) to label microglial cell bodies and anti-CD68 antibody (detected by donkey anti-rat AF555 secondary) to detect CD68 protein. Mounted and cured sections were imaged on Leica Stellaris 8 confocal microscope using a 63x/1.4 NA oil objective (Leica HC PL APO 63x/1,40 OIL CS2). A 3D image stack of 185 μm x 185 μm in XY and approximately 20 μm in the Z dimension, with a voxel size of 72 nm x 72 nm x 299 nm was acquired for each dLGN. The channels were acquired in photon counting mode simultaneously: a 499 nm laser with an emission window of 504-548 nm together with a 553 nm laser with an emission window of 560-650 nm for Iba1-AF488 and CD68-AF555. To eliminate any fluorescence emission bleed-trough between channels, the photon arrival times were gated to 3 - 5 ns for Iba1-AF488 and to 0 - 1.5 ns for Iba1-AF555 channels, respectively. Images acquired were collected as 8-bit datasets, and care was taken to set the laser power, signal accumulation, and pixel dwell time to avoid fluorophore bleaching or detector saturation.

For segmentation of microglia and CD68 signals, we utilized the Surface Creation module with machine learning segmentation of Imaris 10.1.1 (Oxford Instruments). We first trained for several rounds a pixel classifier for each channel then saved the classifiers along with the other settings (smoothing grain size of 145 nm and discarding volumes smaller than 10 voxels) as “Favorite Creation Parameter”. Subsequently, the same Favorite Creation Parameters were applied to all data sets, with the rare instances when microglia segmentation was slightly adjusted via the same pixel classier to better capture some of the cell border areas. Similar to the previous analyses (Figure 5), the segmented signals of CD68-AF555 were converted to binary images (0 or 1 voxels), and the total volume CTB-AF555-positive voxels in each microglia was extracted, converted to fraction of the microglia volume, and plotted. For measurement of relative CD68 intensity, the sum of CD68 signal was divided by the microglia volume. Only the microglia whose body and major processes were confined within the 3D stack were included in the analysis. All staining, imaging and analysis parameters were identically applied to all sections. Two sections were analyzed for each animal. Data from the same genotype and were pooled together.

**Characterization of *App/Ps1* transgenic mice**

To detect Aβ deposition (Figure S13A), brains of 6 month-old female *App/Ps1*^-/-^ mice, 5 month-old and 6 month-old male and female *App/Ps1*^+/-^ mice, and 6 month-old female *Casp3*^-/-^; *App/Ps1*^+/-^ mice were harvested, sectioned, and stained with Thioflavin S as described above. Stained sections were imaged on a Leica Stellaris 8 microscope using a 10x/0.4 NA dry objective (Leica HC PL Apo CS2 10x/0.4 dry). A 460 nm laser line was used for excitation and a window of 469 nm - 600 nm was used for emission. Tile images (pixel size: 284nm x 284 nm) were collected and merged into one image spanning an entire coronal section in Las X 4.6 Navigator. To quantify the level of amyloid deposition in 6 month-old female *App/Ps1*^+/-^ and *Casp3*^-/-^; *App/Ps1*^+/-^ mice (Figure S13B), the number of Thioflavin S-positive plaques was manually counted in a total of 3 sections per animal and averaged.

To investigate the role of caspase-3 in Aβ-induced synapse loss (Figure 7), *Casp3*^-/-^ mice were bred to *App/Ps1*^+/-^ mice to create *Casp3*^-/-^; *App/Ps1*^-/-^ and *Casp3*^-/-^; *App/Ps1*^+/-^ mice. Brains of 6 month-old female *App/Ps1*^-/-^ and *App/Ps1*^+/-^ littermates as well as *Casp3*^-/-^; *App/Ps1*^-/-^ and *Casp3*^-/-^; *App/Ps1*^+/-^ littermates were harvested, sectioned, and stained with anti-SV2 antibody (conjugated to AF546) and anti-Homer1 antibody (using goat anti-rabbit AF647 as the secondary) to label presynaptic and postsynaptic compartments, respectively. Stained sections were imaged on a Leica Stellaris 8 confocal microscope using a 63x/1.4 NA oil objective (Leica HC PL APO 63x/1,40 OIL CS2). To acquire a representative dataset, we focused on the molecular layer of the dentate gyrus of the hippocampus from both the left and right side of the brain and sampled 3 regions from each side, as shown in Figure S14. 3D-stacks (61.5 μm x 61.5 μm x 20 μm) were acquired with a voxel size of 64 nm x 64 nm x 298 nm. 561 nm and 638 nm laser lines were used for excitation, 555 nm – 635 nm and 642 nm – 792 nm windows were used for emission reading, and appropriate notch filters were used for blocking the exciting laser lines. The channels were acquired as 8-bit data sets with detectors set in counting mode, and the lasers power, signal accumulation, and pixel dwell time set at levels that avoided detector saturation. To measure synapse number per unit volume, we first trimmed the raw 3D stacks to a stack of approximately 3.5 μm thickness on the side closest to the coverslip (to avoid any signal depletion the middle part of the section due to antibody penetration issues) and cropped out regions where large gaps (corresponding to blood vessels) devoid of Sv2 or Homer1 staining were present. The punctate signals of SV2 and Homer1 were fit in Imaris 10.1 (Oxford Instruments) to ellipsoids (500 nm in XY and 850 nm in Z for SV2; 400 nm in XY and 800 nm in Z for Homer1) with background subtraction (“Spot Creation”). Then we designated as “synapses” all Homer1 ellipsoids that were within 300 nm of a SV2 ellipsoid (center-to-center distance). For each animal, the total number of synapses in all 3D-stacks analyzed was divided by the total volume of all stacks analyzed and plotted as a single value.

To detect Aβ-induced microglia activation (Figure S16), brains of 6 month-old female *App/Ps1*^-/-^, *App/Ps1*^+/-^, and *Casp3*^-/-^; *App/Ps1*^+/-^ mice were harvested, sectioned, and stained with anti-Iba1 antibody (using goat anti-rabbit AF647plus as the secondary) and anti-Cd68 (using goat anti-rat AF555plus as the secondary) as labels of microglia cell body and microglial activation, respectively. Stained sections were imaged on a Leica Stellaris 8 confocal microscope using a 20x/0.7 NA water objective (Leica HC PL APO 20x/0,7 IMM CORR CS2). Tile images spanning the hippocampus were acquired at voxel sizes of 142 nm x 142 nm x 300 nm. A 548 nm laser line and a 554 nm – 634 nm emission window were used for the CD68-AF555 channel, and a 653 laser line and a 663 nm – 748 nm emission window were used for the Iba1-AF647 channel. Microgliosis could be detected as clusters of microglia with stubby processes and strong intracellular Cd68 signals (Figure S16).

To quantify Aβ-induced caspase-3 activity (Figure S17), brains of 4 month-old, 5 month-old and 6 month-old female *App/Ps1*^-/-^ and *App/Ps1*^+/-^ littermates were harvested, sectioned, and stained with anti-cleaved caspase-3 antibody (using goat anti-rabbit AF555plus as the secondary). Stained sections were imaged on a Leica Stellaris 8 confocal microscope using a 63x/1.4 NA oil objective (Leica HC PL APO 63x/1,40 OIL CS2). We sampled at least 2 regions of interest from the molecular layer of the dentate gyrus of the hippocampus on each side of the brain. 3D-stacks (92.26 μm x 92.26 μm x 22 μm) were acquired with a voxel size of 81 nm x 81 nm x 299 nm. A 561 laser line was used for excitation and an emission window of 550 nm – 728 nm was used for the active caspase-3 channel. To measure the number of caspase 3 objects in the dataset, we first trimmed the 3D stacks to a volume where the signal is uniformly distributed (i.e. in focus) and segmented the objects using the “Surface creation” analysis in Imaris 10.1 (Oxford Instruments). The dataset was first smoothed by a two-pixel filter, then a ball size of 400 nm was used to find the objects with background subtraction. Only objects bigger than 10 and smaller than 2000 voxels were kept. Finally, the number of caspase 3 objects was divided by the volume of the 3D stack analyzed and reported per brain side (left or right) for each animal.

To measure the relative density of granule cells in the dentate gyrus (Figure S16), brains of 6 month-old female *App/Ps1*^-/-^, *App/Ps1*^+/-^, *Casp3*^-/-^; *App/Ps1*^-/-^, and *Casp3*^-/-^; *App/Ps1*^+/-^ mice were harvested, sectioned, and stained with anti-NeuN antibody (using goat anti-mouse AF594plus as the secondary) to label neuronal nuclei. Stained sections were imaged on a Leica Stellaris 8 confocal microscope using a 20x/0.7 NA water objective (Leica HC PL APO 20x/0,7 IMM CORR CS2). 3D stacks of the dentate gyrus (581 μm x 581 μm x 30 μm) were acquired at voxel sizes of 160 nm x 160 nm x 1000 nm for each section and side (left and right) using a 594 nm laser and an emission window of 597 nm to 750 nm. The channels were acquired as 8-bit images with the detector set in counting mode, and the lasers power, signal accumulation, and pixel dwell time set at levels that avoided detector saturation. To count the number of nuclei, images were first rotated to position vertically the cortical side of the dentate gyrus. A maximum projection of the brightest two Z-slices was extracted from each 3D stack, and NeuN+ nuclei were manually counted within a rectangular region of interest (89 μm x 150 μm) that was positioned on the granule cell layer at 300 μm away from the tip of the gyrus.
